# Cyclin A triggers Mitosis either via Greatwall or Cyclin B

**DOI:** 10.1101/501684

**Authors:** Nadia Hégarat, Adrijana Crncec, Maria F. Suarez Peredoa Rodriguez, Fabio Echegaray Iturra, Yan Gu, Paul F. Lang, Alexis R. Barr, Chris Bakal, Masato T. Kanemaki, Angus I. Lamond, Bela Novak, Tony Ly, Helfrid Hochegger

**Affiliations:** Genome Damage and Stability Centre, School of Life Sciences, University of Sussex, Brighton BN19RQ, UK; Department of Biochemistry, University of Oxford, South Park Road, Oxford OX13QU, UK; MRC London Institute of Medical Science, Imperial College, London W12 0NN, UK; The Institute of Cancer Research, London SW3 6JB, UK; National Institute of Genetics, Research Organization of Information and Systems (ROIS), and Department of Genetics, SOKENDAI (The Graduate University of Advanced Studies), Yata 1111, Mishima, Shizuoka 411-8540, Japan; Centre for Gene Regulation and Expression, School of Life Sciences, University of Dundee, Dundee DD1 5EH, UK; Wellcome Trust Centre for Cell Biology, University of Edinburgh, Edinburgh EH9 3BF, UK

## Abstract

Two mitotic Cyclins, A and B, exist in higher eukaryotes, but their specialised functions in mitosis are poorly understood. Using degron tags we analyse how acute depletion of these proteins affects mitosis. Loss of Cyclin A in G2-phase prevents the initial activation of Cdk1. Cells lacking Cyclin B can enter mitosis and phosphorylate most mitotic proteins, because of parallel PP2A:B55 phosphatase inactivation by Greatwall kinase. The final barrier to mitotic establishment corresponds to nuclear envelope breakdown that requires a decisive shift in the balance of Cdk1 and PP2A:B55 activity. Beyond this point Cyclin B/Cdk1 is essential to phosphorylate a distinct subset mitotic Cdk1 substrates that are essential to complete cell division. Our results identify how Cyclin A, B and Greatwall coordinate mitotic progression by increasing levels of Cdk1-dependent substrate phosphorylation.

## Main Text

Cdk1 phosphorylates over 1000 proteins^1,2^ within the brief 20-30 minute window of mitotic entry, triggering centrosome separation and chromosome condensation in prophase, followed by nuclear envelope breakdown (NEBD) and mitotic spindle formation in prometaphase, and the alignment of bi-oriented sister chromatids at the metaphase plate^3^. Binding of a Cyclin partner is critical for allosteric activation of CDKs. Two families of mitotic Cyclins, termed A and B, work with Cdk1 to orchestrate mitotic entry in higher eukaryotes^4,5^. Despite the central importance of these proteins for cell cycle control, the functional specialisation of mammalian A and B-type Cyclins remains unclear. Following the depletion of maternal pools of early embryonic Cyclin A1 and B3, somatic mammalian cells express one A-type Cyclin, A2, and two B-type cyclins, B1 and B2.

Genetic depletion in mice suggests an essential role for Cyclin A2 for development, but not for embryonic fibroblast proliferation^6^. Depletion of Human Cyclin A2 by siRNA delays mitotic entry, and this is further enhanced by co-depletion of Cyclin B1^7–9^. Likewise, work in mammalian cell extracts documented how Cyclin A synergises with Cyclin B to control the mitotic entry threshold at the level for Cdk1 activation^10^. A mechanism involving Plk1 activation has been suggested^11,12^, although an essential role of Plk1 in the G2/M transition remains contentious^13^. Likewise, work in mammalian cell extracts documented how Cyclin A synergises with Cyclin B to control the mitotic entry threshold at the level for Cdk1 activation^10^. A confounding factor in the genetic analysis of Cyclin A2 has been its dual role in S-phase and mitosis^14^, making it difficult to directly investigate G2 specific defects.

Murine Cyclin B1 is essential for development^15^ and critical for mitotic entry in early mouse embryos^16^. Conversely, mice lacking Cyclin B2 live healthily, without apparent defects^15^. These results stand in stark contrast to observations from experiments involving siRNA depletion of B-type Cyclins in human cancer cell lines that show surprisingly mild mitotic entry defects^9,17^ The observed unperturbed G2/M transition in Cyclin B depleted human cells could be explained by a synergy between A and B-type cyclins in mitotic entry^9^.

In parallel to mitotic Cyclin/Cdk1 activation, the inactivation of the Cdk1 counteracting phosphatase PP2A:B55 by Greatwall kinase (Gwl) via its substrates Ensa and ARPP19 also plays a critical role in mitotic entry in Xenopus egg extracts^18–20^. However, human and mouse cells lacking Greatwall kinase readily enter mitosis^21,22^, but fail to undergo accurate sister chromatid segregation and cell division^23^. Thus, it appears that human cells can enter mitosis in the absence of Cyclin B or Greatwall, but it remains unclear to what extent the kinase activation and phosphatase inactivation pathways compensate for each other. Moreover, the precise G2 specific role of Cyclin A in this regulatory network is still an open question. In this study, we have taken advantage of an optimised degron tagging approach to address these questions.

Rapid, precise, and uniform depletion is critical to investigate the functions of mitotic Cyclins within a single cell cycle. We have recently reported that a combination of AID-^24^ and SMASh-^25^ tags results in highly efficient induced degradation^26^, and applied this strategy to comprehensively analyse the functions of Cyclin A2 and B1 in a non-transformed, h-TERT immortalised human epithelial cell line, RPE-1 (Figure S1). To further reduce the levels of B-type Cyclins in these cells we also generated a CRISPR based knock out of the CCNB2 gene encoding Cyclin B2 in the Cyclin B1 degron cells, and verified absence of Cyclin B2 by western blotting (Figure S1D). The homozygous Cyclin A2 and B1 double degron mAID-SMASh tag fusions (A2^dd^, B1^dd^) allowed us to compare the depletion efficiency of the single and combined degron systems in these cells (Figure S2). These results demonstrate that the individual degron tags were not sufficient to fully deplete the tagged Cyclins, likely due to their rapid synthesis rates. Activation of both degradation systems using the “DIA” cocktail (Dox/IAA for mAID and ASV for SMASh, see methods), on the other hand, was highly effective to trigger rapid degradation of either Cyclin that became undetectable after 2-4 hours of treatment. Quantification of this depletion experiment (Figure 1A and B) shows 50% depletion within the first hour and an ultimate depletion to less than 5% of control protein levels within 2-4 hours.

**Figure 1:**
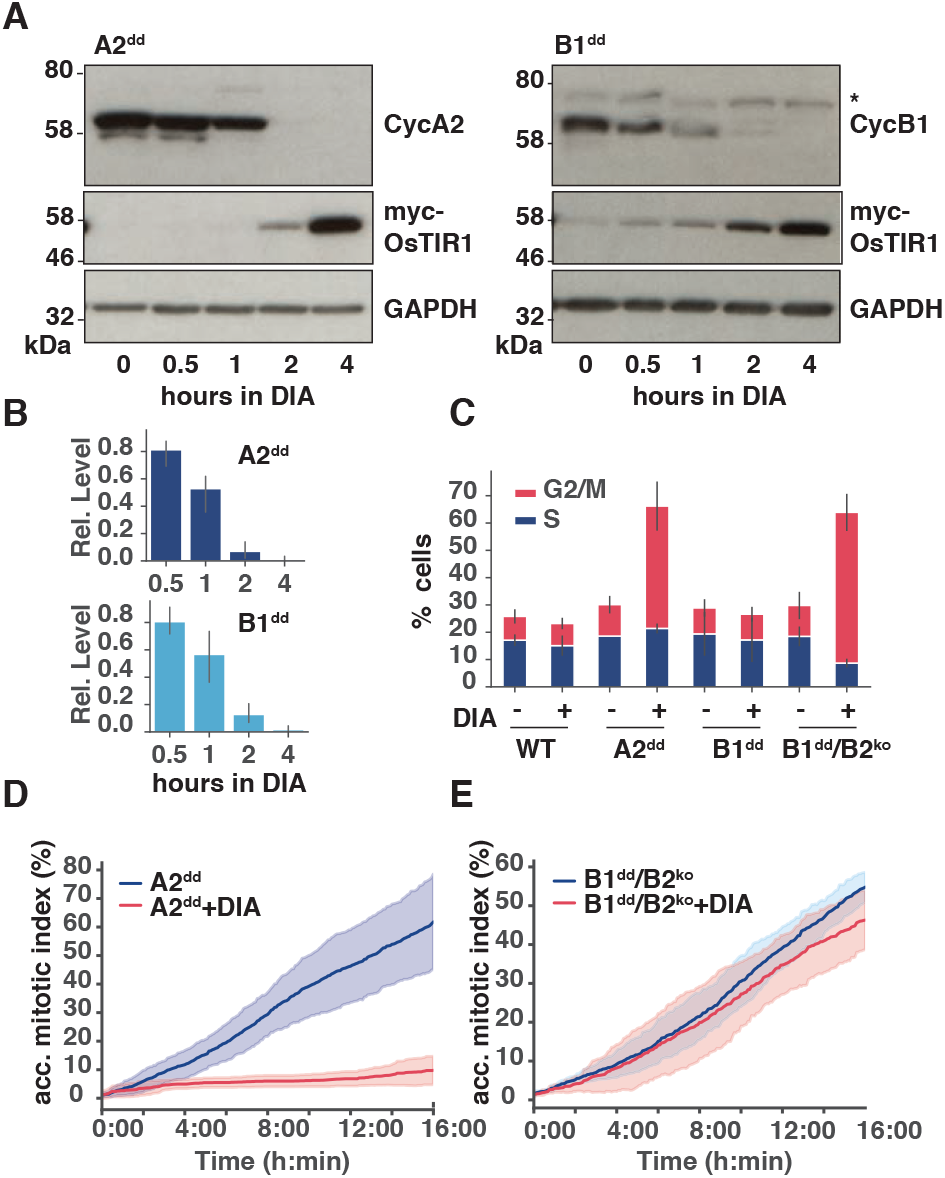
Cyclin A, but not B is required for mitotic entry in RPE-1 cells. (**A**) Representative immuno-blots and (B) quantitation showing time course of induced degradation of double degron tagged Cyclin A and Cyclin B (A2^dd^, B1^dd^). * indicates cross reacting band. (n=3 experiments, s.d. indicated by error bars). (**C**) Cell cycle phase frequencies measured by flow cytometry in indicated cell lines (see Figure S3A for raw FACS data) 24 hours after DIA treatment, (n=3 experiments, s.d. indicated by error bars). (**D**) Kinetics of mitotic entry as measured by time-lapse microscopy in A2^dd^ cells following mock or DIA treatment. Cells were imaged for 16h with 5min intervals. Curves display accumulative mitotic index (Data from three repeats, n>500 cells per condition, stdv indicated by shaded area). (**E**) Kinetics of mitotic entry as in (D) in B1^dd^/B2^ko^ cells

Using this depletion system, we further analysed A2^dd^, B1^dd^ and B1^dd^/B2^ko^ cells by measuring the distribution of cell cycle phases in asynchronous cells, 24 hours after DIA treatment (Figure 1C and Figure S3A). Loss of Cyclin A2 caused an accumulation of the 4N population, but depletion of Cyclin B1 had little effect on the cell cycle phase distribution of RPE-1 cells. Furthermore, DIA treatment abolished cell proliferation in A2^dd^ cells, but only had a mild effect in Cyclin B1^dd^ cells as judged by a long-term proliferation assay (Figure S3B). RPE-1 cells lacking Cyclin B1 relied on Cyclin B2 for proliferation, because B1^dd^/B2^ko^ cells showed an accumulation of 4N cells and decrease in proliferation following DIA treatment (Figure 1C and S3). We next measured mitotic entry dynamics in cells lacking either Cyclin A2 or both B1 and B2 (Figure 1D and E, Figure S4). DIA treated A2^dd^ cells ceased to enter mitosis rapidly following drug addition, while lack of Cyclin B1 and B2 did not measurably affect the dynamics of mitotic entry. However, these cells had severe defects in mitotic progression and cell division (see below). We also tested, if Cyclin B3 could account for mitotic entry observed in the absence of Cyclin B1 and B2. However, we could not detect Cyclin B3 protein in these cells by immuno-blotting despite the presence of the mRNA of this Cyclin. Moreover, siRNA depletion of the Cyclin B3 mRNA in combination with DIA treatment of B1^dd^/B2^ko^ cells did not appear to reduce the number of mitotic cells (Figure S3C and D). Taken together, these results suggest that Cyclin A2 is essential to initiate mitosis and that B-type Cyclins are not required for mitotic entry.

Previous reports implicated Cyclin A2 in triggering M-phase via Cdk1 activation^7,9^, but could not differentiate between knock-on effects of DNA replication problems and G2 specific functions of Cyclin A2. We took advantage of the A2^dd^ cells to address this question by performing G2 specific depletion of Cyclin A2. For this purpose, we used a PCNA-mRuby fusion^27^ to identify cells that were in G2 phase at the time of DIA treatment (Figure 2A). This assay revealed that DIA addition in G2 phase effectively blocked further progression into mitosis (Figure 2B), suggesting that either the presence, or new synthesis of Cyclin A in this cell cycle phase is required for mitotic entry. To further support this notion, we performed EdU pulse labelling at the time of DIA treatment, followed by fixation and CENP-F immuno-fluorescence five hours later to mark cells that were in G2 phase (CENP-F positive/EdU negative) when Cyclin A2 degradation began (Figure S5). This resulted in an accumulation of cells that had been in G2 phase at the time of DIA treatment, further supporting a G2-specific role for Cyclin A2. Inhibition of Wee1, a kinase that inhibits Cdk1 in interphase, was sufficient to overcome this block and resulted in resumption of mitotic entry with kinetics comparable to the controls (Figure 2C). Once the G2 block was overcome by Wee1 inhibition, cells lacking Cyclin A could form a metaphase spindle and underwent chromosome segregation and cell division (Figure 2D and E). Thus, Cyclin A does not appear to play an essential role in mitosis once Cdk1 is activated.

**Figure 2:**
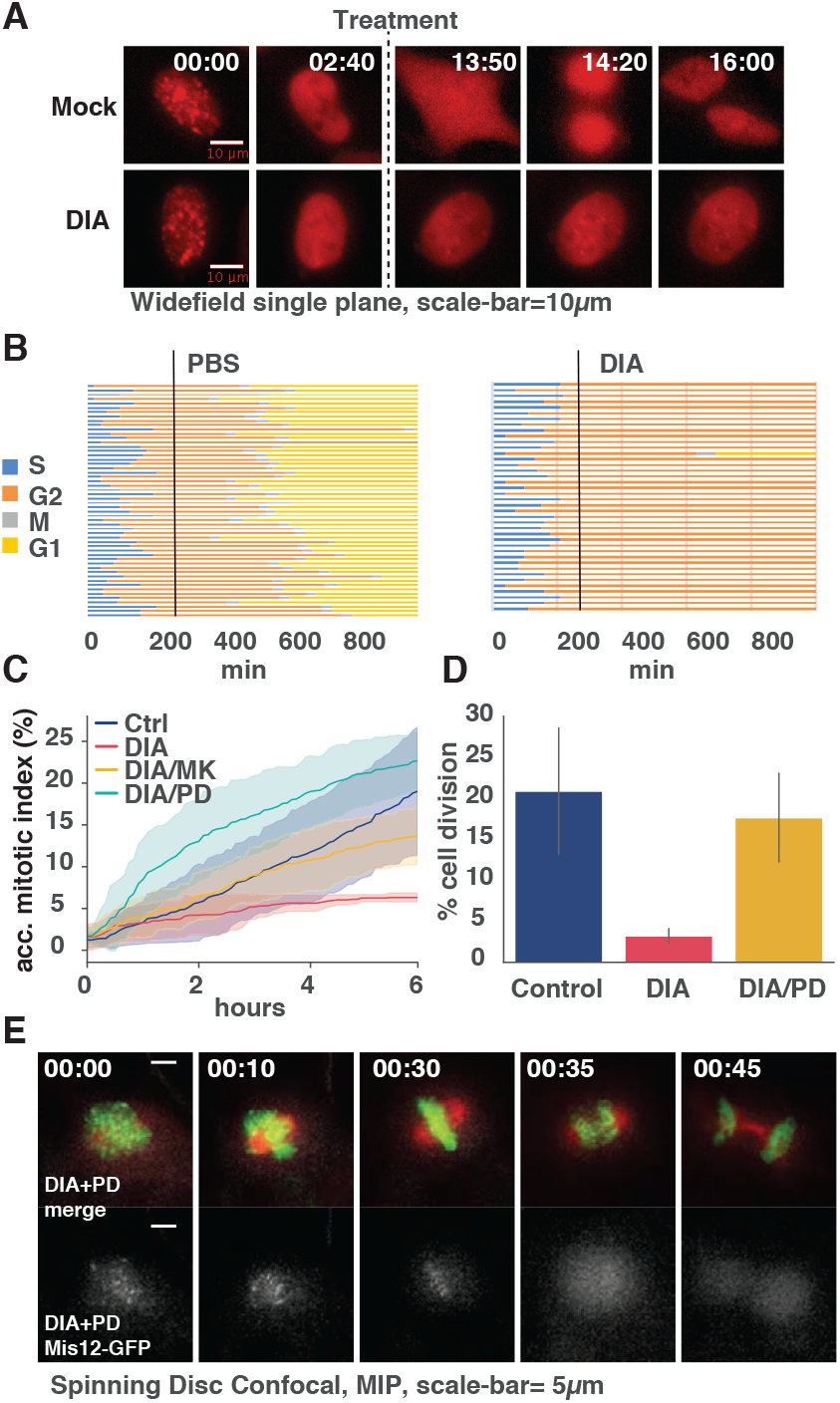
Cyclin A2 is essential to trigger mitotic entry in G2 phase but not required for further mitotic progression. **(A)** Images of time-lapse experiment with PCNA-mRuby tagged A2^dd^ cells. The imaging sequence was started at the time of Doxycyclin addition, and degron activity was triggered three hours later by addition of IAA and Asv or PBS (indicated by dashed line). Time is shown as hh:min. **(B)** Single cell analysis of 40 cells per condition treated as described in (A) **(C)** Kinetics of mitotic entry, measured by time-lapse microscopy in A2^dd^ cells following mock or DIA treatment and addition of Wee1 inhibitors MK1775 (1μM) or PD-166285 (0.5μM). Cells were imaged for 6 hours in 5min intervals. (Data from 3 repeats, n> 500 cells per condition, s.d. indicated by shaded area). **(D)** Frequency of successfully completed cell division in A2^dd^ cells treated with DIA and PD-166285. Data represent mean value of three independent experiments (n>50 per repeat, error bars show stdv). **(E)** Images from time lapse sequence of mitosis showing cell division after addition of PD 188285 in DIA treated A2^dd^ expressing FusionRed-H2B (upper panel, green) and Mis12-GFP (lower panel, white) and labelled with SiR-tubulin (upper panel, red).

We subsequently analysed the phenotype of cells lacking B-type Cyclins in closer detail. DIA treated B1^dd^/B2^ko^ cells readily entered mitosis but failed to undergo sister-chromatid segregation and cytokinesis (Figure 3A and S6). Indeed, in more than 150 mitotic events observed overall, we did not record a single successful example of normal mitotic exit in cells lacking Cyclin B1 and B2, while cells lacking either B1 or B2 alone did not have any apparent mitotic defects. Even in cases where DIA-treated B1^dd^/B2^ko^ cells initiated segregation and cytokinesis, all cells ultimately failed cell division, resulting in a single bi-nucleated daughter cell. A fraction of these cells did not transit prophase and exited mitosis without undergoing NEBD. Moreover, we observed a delay in mitotic progression following DIA treatment (Figure 3B). To analyse this phenotype with improved resolution we recorded mitotic progression in B1^dd^/B2^ko^ cells that stably expressed RFP-Histone H2B and the kinetochore component Mis12-GFP. Figure 3C shows a typical example of mitotic progression in cells with or without B-type cyclins (see also Videos 01 and 02). While cells lacking cyclin B can clearly initiate chromosome condensation, NEBD and spindle formation they fail to segregate their sister chromatids and exit mitosis with a single fragmented nucleus.

**Figure 3.**
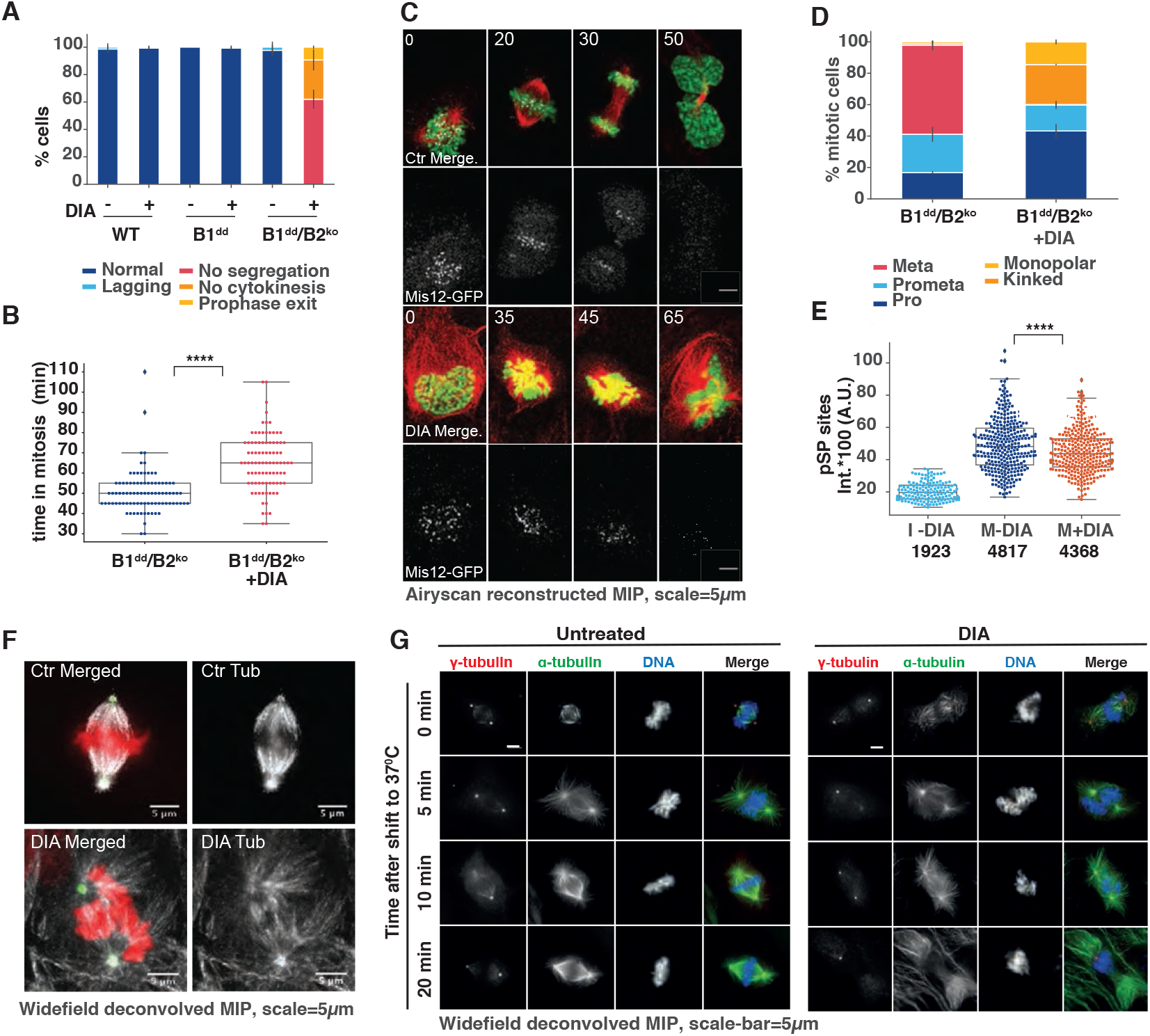
Cyclin B is essential for sister chromatid bi-orientation and segregation. **(A)** Frequency of mitotic phenotypes observed in indicated cell lines with or without DIA treatment (3 repeats, n>50 per repeat, error-bars indicate s.d.) **(B)** Mitotic duration comparing mock-treated and DIA-treated B1^dd^/B2^ko^ cells. Time of Entry and Exit was defined by chromosomes condensation and decondensation. (3 repeats, n>50 per repeat, error bars indicate s.d.) **(C)** Representative images from live cell imaging of SiR-Tubulin labelled (red) mitotic B1^dd^/B2^ko^ cells expressing FusionRed-H2B (green), Mis12-GFP (white); time in mins. **(D)** Characterisation of mitotic spindle in Pro-tame/Apcin (P/A) arrested B1^dd^/B2^ko^ cells with or without DIA treatment (3 repeats, n>50 per repeat, error bars indicate s.d.). For cell synchronisation (P/A synchronisation) we released from 24 hour Thymidine block into DIA- or PBS-containing medium and added (P/A) 10 hours after release. The APC/C inhibition lasted for two hours, and was followed by fixation and immuno-fluo-rescence analysis. **(E)** Intensities of anti-(pSP) antibody staining in cells treated as in (D) (3 repeats, n>50 per repeat). Key to Legend: I-DIA = control interphase cells, M-DIA, M+DIA = control and DIA treated mitotic cells. Numbers below panel indicate median values **(F)** Representative images of mitotic spindles in control- and DIA-treated B1^dd^/B2^ko^ cells as in (D). (Tubulin (white), Pericentrin (green), DAPI (red)). **(G)** Immuno-fluorescence images from mitotic Ctr and DIA treated B1^dd^/B2^ko^ cells stained with anti-tubulin (green) and anti-gamma-tubulin (red) antibodies and DAPI (blue). Cells were exposed to ice cold medium and either fixed immediately after cold exposure, or incubated for 5,10, or 20 minutes at 37C before fixation.

A simple explanation for this failed mitotic exit could be the early degradation of Cyclin A2 by the APC/C that is independent of the spindle assembly checkpoint. However, we observed a distinct mitotic defect in cells lacking Cyclin B1/2 (Figure 3D-F) even in the presence of a APC/C inhibitor cocktail^28^. Thus, mitotic cells expressing only Cyclin A2, but lacking B-type Cyclins assemble a bipolar, but often malformed spindle, and fail to align their chromosomes on the metaphase plate. Overall, we observed an enrichment in prophase, and prometaphase (or defective metaphase) after DIA treatment and APC/C inhibition, but failed to identify DIA treated cells with a clear metaphase alignment (Figure 3F). Exposure to ice cold medium revealed intact K-fibres in DIA treated B1^dd^/B2^ko^ cells, and re-exposure to warm medium resulted in the reformation of a disorganised spindle with extended astral microtubules (Figure 3G). We also used the centromere/kinetochore markers CenpA and CenpB to asses bi-orientation and segregation of sister chromatids (Figure S7). The markedly reduced distance between sister centromeres following DIA treatment suggests lack of tension. Moreover, we frequently observed unseparated centromere pairs in cells that were exiting mitosis, as judged by attempted furrow formation, indicating that Separase activation and/or MT force generation was impaired in these cells. These distinct phenotypes suggest that, while Cyclin B is required for specific regulatory events to coordinate the metaphase/anaphase transition, it is not essential for the establishment of the mitotic state. Accordingly, we observed only a small decrease in the level of Cdk substrate phosphorylation following DIA treatment as judged by quantitative immuno-fluorescence (Figure 3E).

To further quantity the contribution of Cyclin B to mitotic phosphorylation events and identify Cyclin B specific substrates, we compared the phospho-proteome in control and DIA treated B1^dd^/B2^ko^ cells (Figure 4A, S8). We quantified differences in phosphorylation in 5,760 phosphorylation-sites (Supp. Table 1), including 3,192 that have [S/T]P motif in the peptide sequence detected, and identified 360 sites in 263 substrates that were reduced by more than 30%. Thus, Cyclin B depletion causes a significant but relatively minor effect on the majority of mitotic phosphorylations, but causes a more dramatic reduction in phosphorylation of only ~5% of the phospho-proteome, and ~8.4% of the proline-directed sites (271 / 3,192). These substrates were predominantly cytoskeletal and chromatin associated proteins (Figure 4B). A more detailed network analysis (Figure S9) identified a variety of mitosis associated processes that involve Cyclin B specific substrates, including chromosome structure, MT dynamics and cytokinesis.

**Figure 4.**
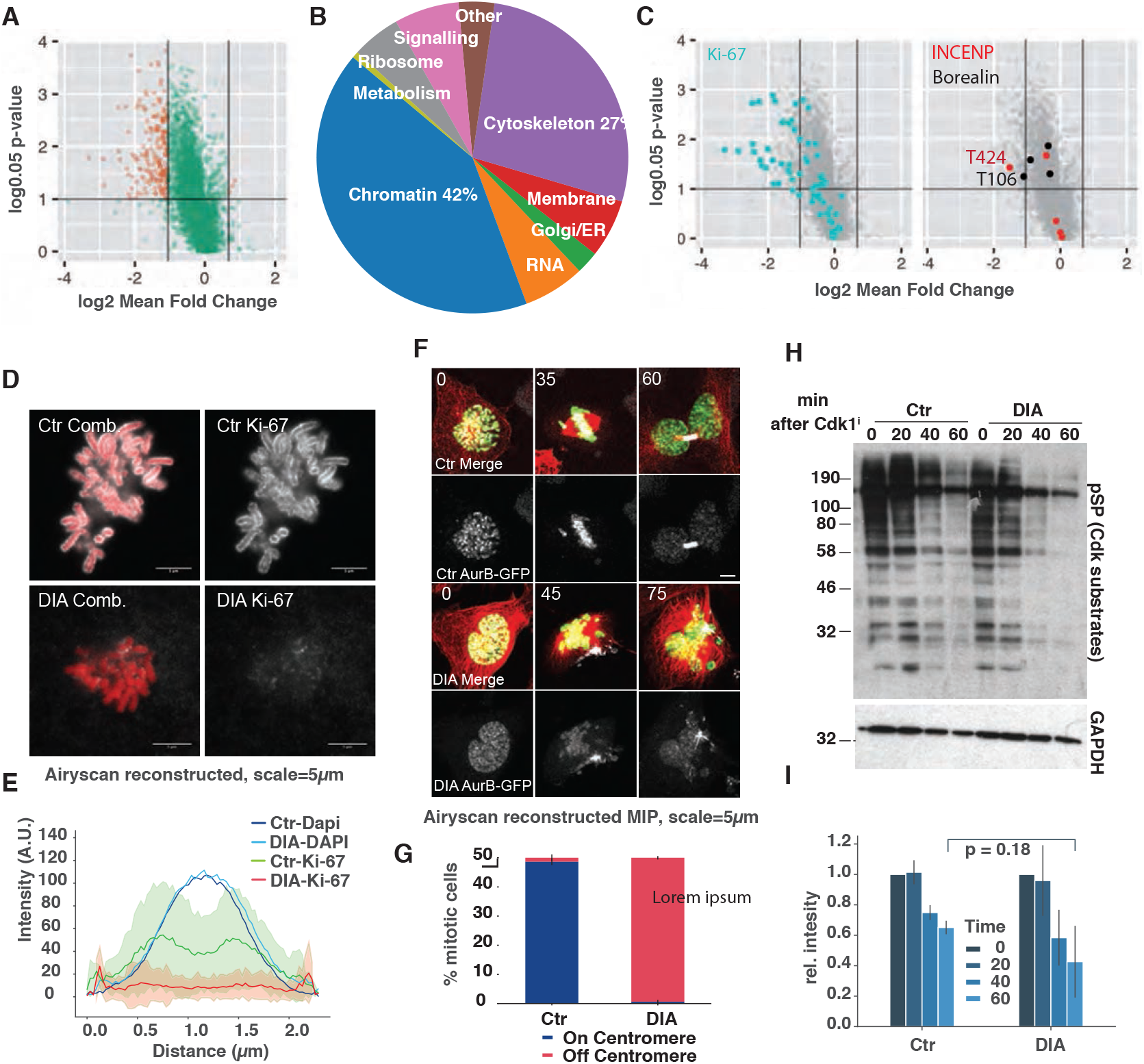
Cyclin B is not essential for the bulk of mitotic substrate phosphorylation, but only required for a specific subset of substrates. **(A)** Quantification of Cyclin B specific substrates by phosphoproteomics. Phosphorylation sites deemed significantly changing are shown in red (p < 0.05, 1.96 * s.d. fold change cutoffs see Methods). **(B)** Gene ontology analysis of Cyclin B-dependent substrates. **(C)** Quantification of changes in KI67 and INCENP/Borealin phosphorylation after Cyclin B depletion **(D)** Representative images of Ki-67 immunofluorescent staining of chromosome spreads from control (Ctr) and DIA treated P/A synchronised B1^dd^/B2^ko^ cells **(E)** Quantification of Ki-67 staining intensity (images from (I) of cross sections through chromosomes (Data from three repeats, n=50, shaded area indicates s.d.). **(F)** Representative images from live cell imaging of DIA-treated SiR-tubulin labelled (red) B1^dd^/B2^ko^ cells expressing FusionRed-H2B (green) and AurB-GFP (white); time in mins. **(G)** Frequencies of aberrant AurB localisation in P/A synchronised B1^dd^/B2^ko^ cells. (Bars indicate mean of three repeats, n=50 cells per repeat). **(H)** Dynamics of mitotic substrate dephosphorylation triggered by Cdk1 inhibition in control and DIA treated B1^dd^/B2^ko^ cells. The cells were enriched in mitosis by thymidine release and Apcin/ProTame treatment and treated with 1μM Flavo-puridinol to trigger mitotic exit. Dephopshorylation was measured by probing the immune-blots with anti Cdk1 substrate (pSP) antibody. **(I)** Three repeats of the experiment in (G) were quantified and corrected based on the corresponding GAPDH intensity values. The bar plots show the eman of three experimenst and the standard deviation is indicated by error bars.

We validated this data set experimentally by analysing the localisation of a group of Cyclin B specific targets identified in the screen. For this purpose, we chose Ki-67 as well the Chromosome Passenger complex (CPC) members INCENP and Borealin. Each of these proteins shows a significant loss of phosphorylation after DIA treatment in the mass spectrometry analysis (Figure 4C, Supp. Table 2). Ki-67 acts as a surfactant on condensed mitotic chromosomes^29^ and is heavily phosphorylated by Cdk1 in mitosis^30^. Loss of Cyclin B following DIA treatment resulted in a loss of Ki-67 from the chromosome periphery (Figure 4D and E), correlating with the marked decrease in multiple phosphorylations in Ki-67 in DIA treated B1^dd^/B2^ko^ cells. Likewise, INCENP and Borealin have previously been identified as mitotic targets of Cdk1^1,2,31,32^, but the Cyclin B specific phospho-sites that we have identified here (T424 in INCENP and T106 in Borealin) have, to our knowledge, not previously been related to CPC localisation and function. DIA treatment resulted in premature displacement of a key CPC subunit, AurB-GFP, from the centromeres (Figure 4F, Video 03 and 04) in the large majority of DIA treated B1^dd^/B2^ko^ cells, even when cells were arrested in mitosis by APC/C inhibition (Figure 4G). CPC mis-regulation also correlates with the significant decrease in Histone H3 S10 phosphorylation, a target of AurB kinase activity that we observed (Supp. Table 2). Likewise, the reduction of phosphorylation of CdcA2, Top2B and Tpx2 following DIA treatment correlated with changes mitotic localisation (Figure S10), further validating the significance of our proteomic data set.

The question remains, how cells can cope with loss of the major mitotic kinase Cdk1/Cyclin B, and still readily phosphorylate most of the mitotic phospho-proteome, including many bona-fide Cdk sites? A simple answer to this question could be that Cyclin A has two avenues to establish the mitotic state: (i) by activating Cyclin B/Cdk1, (ii) by inactivating PP2A:B55 via Greatwall kinase dependant phosphorylation of ARPP19 and Ensa. In this case, the Greatwall pathway should be fully functional, in cells lacking B-type cyclins. This appears to be the case, since Ensa/ARPP19 phosphorylation at S67 (Supp. Table 2), and dephosphorylation dynamics of mitotic Cdk substrates following Cdk inhibition are not significantly changed in DIA treated B1^dd^/B2^ko^ cells (Figure 4H and I, and S11A).

If the inhibition of PP2A:B55 by phosphorylated Ensa/ARPP19 is responsible for establishing mitosis in the absence of Cyclin B, simultaneous depletion of B-type Cyclins and the Greatwall pathway should ultimately prevent cells from entering mitosis. To test this hypothesis, we depleted Ensa and ARPP19 from B1^dd^/B2^ko^ cells (Figure S11A) and monitored mitotic progression by time-lapse microscopy (Figure 5A, Videos 05 and 06). Depletion of Ensa/ARPP19 did not prevent mitotic entry, but resulted in sister chromatid misalignment and cytokinesis defects, as previously reported for cells lacking Greatwall^33^. Cyclin B depletion by DIA treatment caused similar mitotic exit phenotypes as described above (Figure 3A-D). Surprisingly, cells with simultaneous depletion of Ensa/ARPP19 and Cyclin B appeared able to enter prophase, as judged by cell rounding, centro-some separation and compaction of the nucleus. However, these cells did not undergo NEBD and often remained in this prophase-like state for a prolonged period of time, before reverting to interphase (Figure 5A, bottom panel, Figure S11C, Video 06).

**Figure 5.**
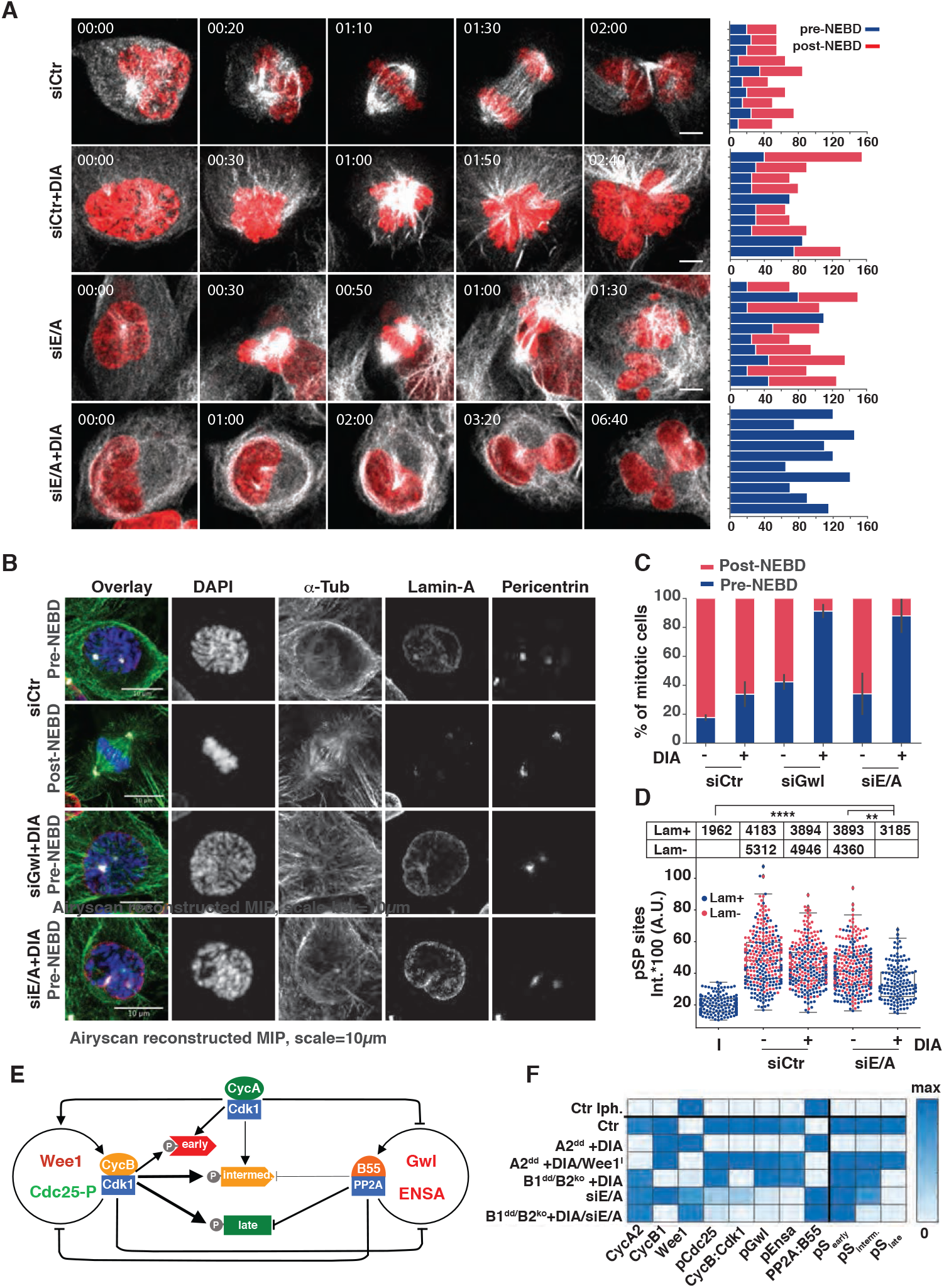
Cells deficient in the Greatwall pathway and Cyclin B arrest in prophase. **(A)** Representative images from live cell imaging of siRNA transfected B1^dd^/B2^ko^ cells (FusionRed-H2B in red, SiR-Tubulin in white). Bar-plot panels on the right show single cell analysis of 10 cells manually scored for length mitosis pre-NEBD and post-NEBD. **(B)** Panels of representative immuno-fluorescence images of siRNA transfected and P/A treated cells treated as indicated. **(C)** Quantification number of mitotic B1^dd^/B2^ko^ cells following siRNA transfection and/or DIA treatment. Frequencies of mitotic cells with intact (blue) and disassembled (red) Lamin A/C staining are plotted (mean value of three repeats, error bars indicate stdv). **(D)** Intensity of pSP-antibody staining in siRNA transfected and DIA treated B1^dd^/B2^ko^ cells. The swarm-blots classify data from mitotic cells with intact (blue) or disassembled (red) Lamin A/C. The table above the panel indicates the median values and selected p-values are indicated. Data are from three repeats, n>50 per repeat. **(E)** Influence diagram of the network controlling the mitotic substrates phosphorylation. The autoactivation of CycB:Cdk1 and PP2A:B55 is controlled by inhibitory phosphorylation and by the Gwl-ENSA pathway, respectively. CycA:Cdk1 helps CycB:Cdk1 auto-activation and inhibits PP2:B55 activity through Gwl phosphorylation. Three different classes of mitotic substrates (early, intermediate and late) are distinguished based on their sensitivity to CycA:Cdk1, CycB:Cdk1 and PP2A:B55. **(F)** Final level of mitotic regulators and Cdk1-substrates in response to cyclin synthesis. The heat map is based on model simulations (see Figure S14) starting with initial conditions without cyclins (Interphase, top row) until high and constant cyclin levels caused by APC/C inhibition.

These data suggest that cells can enter prophase in the absence of Cyclin B and with high PP2A:B55 activity, but fail to move past NEBD. To further substantiate this observation we analysed the mitotic state following Greatwall/Cyclin B and Ensa/ARPP/Cyclin B depletion (Figure S11A, B) in fixed cells by immuno-fluorescence (see representative images in Figure 5B). The mitotic index, as judged by cell rounding, centrosome separation and chromosome condensation, was only mildly reduced in the co-depleted cells (Figure S11D). However, more than 90% of mitotic cells lacking both Cyclin B and Greatwall or Ensa/ARPP19 failed to undergo NEBD as judged by Lamin A/C staining (Figure 5C).

Quantification of various mitotic indicators further supported this observation. Cells lacking both Cyclin B and Ensa/ARPP19 showed levels of Serine-Proline phosphorylation that were clearly increased compared to interphase cells, and only marginally decreased compared to control prophase cells (Figure 5D). Phosphorylation of Lamin A/C S22, a residue that is phosphorylated in late G2/early prophase^34^, was detected at a level comparable to control prophase and mitotic cells (Figure S11E). Likewise, centrosomes were at a stage of separation similar to controls before NEBD (Figure S11F), suggesting that these cells indeed remained in a prophase-like state without progressing past NEBD.

Our results can be summarised by a mathematical model (Figure 5E, F and S12) that suggests a simple ordering mechanism for mitotic entry, based on the increasing requirement for Cdk1-Cyclin activity, and increasing levels of sensitivity to the antagonising phosphatase activity. This fits well with a similar biochemical model for the ordering of mitotic exit^23,35^. Cyclin A in G2 phase is critical to trigger mitosis by counteracting Wee1 and is sufficient to phosphorylate an early set of Cdk1 substrates that cause the initiation of prophase. These substrates are likely to be poor PP2A:B55 targets and therefore require a low threshold of Cdk1:Cyclin activity. In the absence of Greatwall and Cyclin B, cells remain in this prophase state for several hours before reverting back to interphase. This observation is supported by our recent report of a latent stable steady state that can be reached in prophase^36^.

The next step in mitotic establishment, NEBD and transition to prometaphase, requires either Greatwall dependent PP2A:B55 inactivation, or Cyclin B dependent increase in the levels of Cdk1 activity to phosphorylate the bulk of intermediate mitotic substrates. Our finding that Cyclin B and Greatwall act in parallel explains how this regulatory network is designed to allow buffering against fluctuations in kinase and phosphatase activity during mitotic entry. It also explains the lack of entry phenotypes in Greatwall and Cyclin B depleted cells.

Our study defines a final set of late substrates that strictly require Cyclin B dependent Cdk1 activity. These phosphorylation events are critical to initiate anaphase and cell division. We assume that these phospho-sites are highly sensitive to Cdk1 counteracting phosphatases and therefore require higher levels of Cdk1:Cyclin activity. However, it remains to be seen, what ultimately determines this specific requirement for B-type Cyclins. Moreover, future work will need to establish, why Cyclin B cannot compensate for Cyclin A2 as the mitotic trigger. The rapid induced degradation system that we report here will be a helpful tool to further address these questions for a better understanding of mammalian cell cycle control mechanisms.

## Acknowledgements

We thank the members of the Hochegger, Ly, Lamond and Novak labs for supporting work on this study. We thank Tim Hunt and Randy Poon for critical evaluation of this MS, and Viji Draviam for discussing results and sharing reagents. HH was a CRUK senior research fellow (C28206/A14499) and TL is supported by a Sir Henry Dale Fellowship jointly funded by the Wellcome Trust and the Royal Society (ID 206211/Z/17/Z). HH and BN were supported by a BBSRC LoLa grant (ID BB/M00354X/1). We acknowledge the support of the Wolfson Foundation (Grant ref 20440) for funding the Zeiss LSM880 and Operetta CLS microscopes used in this study.

---

**Supplementary Figure S1.**
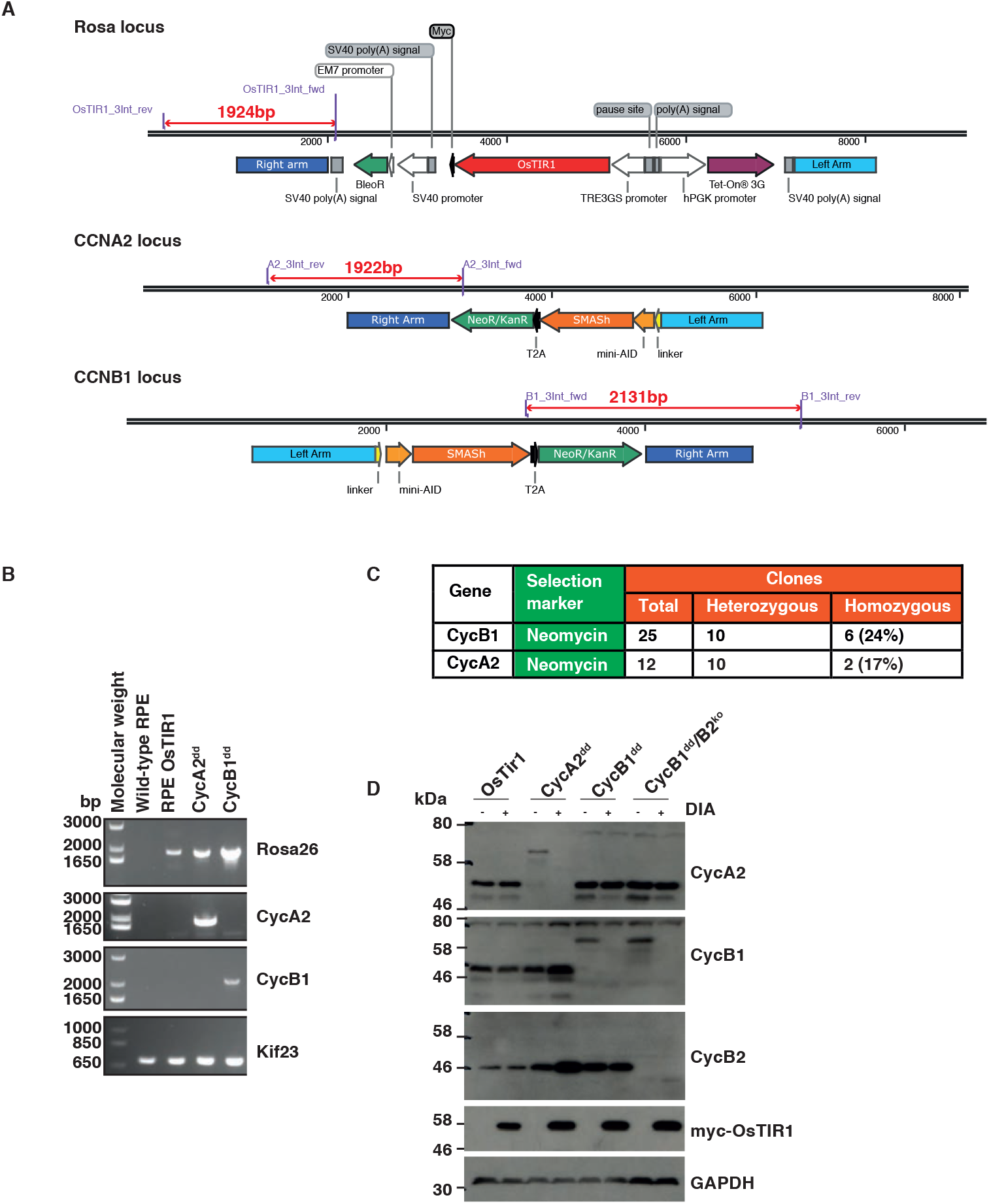
Generation and characterisation of cell lines expressing mAID-SMASh-tagged Cyclin A2 or B1. **(A)** Knock-in strategy for inducible OSTIR1 expression at the human Rosa26 locus, and endogenous Cyclin A2 and B1 protein tagging. We integrated mAID-SMASh-T2A-neomycin in frame and at the C-terminal of the coding sequence of Cyclins A2 and B1. The chosen length for the homologous regions for gene targeting was approximately 1kb. **(B)** To verify gene targeting Primers were designed in the genomic locus and in the cassette (indicated in A) and used to amplify genomic DNA from different cell lines: wild-type RPE-1, RPE-1 expressing OsTIR1, CycA2^dd^, CycB1^dd^, and primers amplifying a region of the Kif23 gene were used as positive control (see Material and Methods section for primer sequences). **(C)** Table showing the success rates of homozygous tag integration after clone screening. **(D)** Confirmation of gene-targeting, Cyclin B2 knock out and induced degradation by immuno-blotting. Indicated cell lines were analysed 24 hours after mock or Dox/IAA/Asv (DIA) treatment using the indicated antibodies to confirm homozygous gene tagging and efficiency of protein degradation.

**Supplementary Figure S2.**
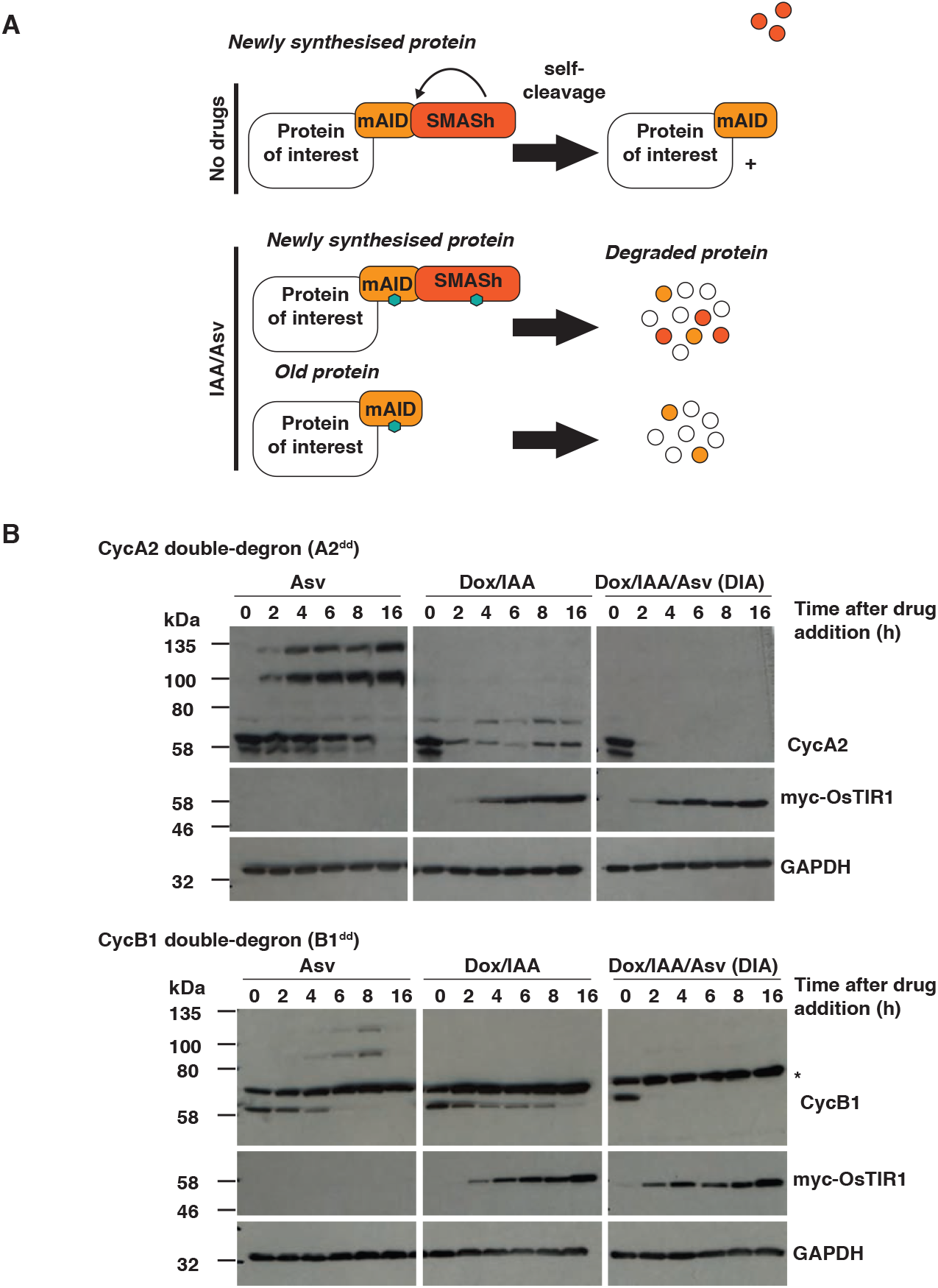
Enhanced protein degradation after knock-in of combined degrons. **(A)** Schematic of protein degradation using AID and SMASh systems. The SMASh tag consists of the self-cleaving viral HCV NS3 protease fused in cis to a constitutive destabilising peptide. The tag detaches itself from the target protein in the off condition. Upon inhibition by the protease inhibitor, asunaprevir (Asv), the tag remains on the target protein and exposes it to the cellular degradation machinery, while the old protein is depleted depending on its specific half-life. The AID tag relies on the expression of the plant F-box protein TIR1 that combines with mammalian Skp1 and Cul1 to form a functional SCF E3 ligase. It is activated to recognise and ubiquitylate the AID degron tag upon stimulation by the hormone Auxin (IAA). In the case of the tag combination newly synthesised proteins are depleted by the two independent degradation systems, while old proteins are destabilised by the mAID tag. **(B)** Comparison of protein degradation efficiency using different degrons. Cells were incubated for inicated time periods with Asv or DOX/IAA or both (DIA) then collected for western blot analysis. Doxycyclin was added 2 hours before addition of IAA or ASV/IAA. To determine protein degradation, either cyclin A2 or cyclin B1 were probed as indicated. Anti-myc antibody was used to check OsTIR1-myc expression. Asteriks indicates non-specific protein cross reacting with the Cyclin B1 antibody.

**Figure S3.**
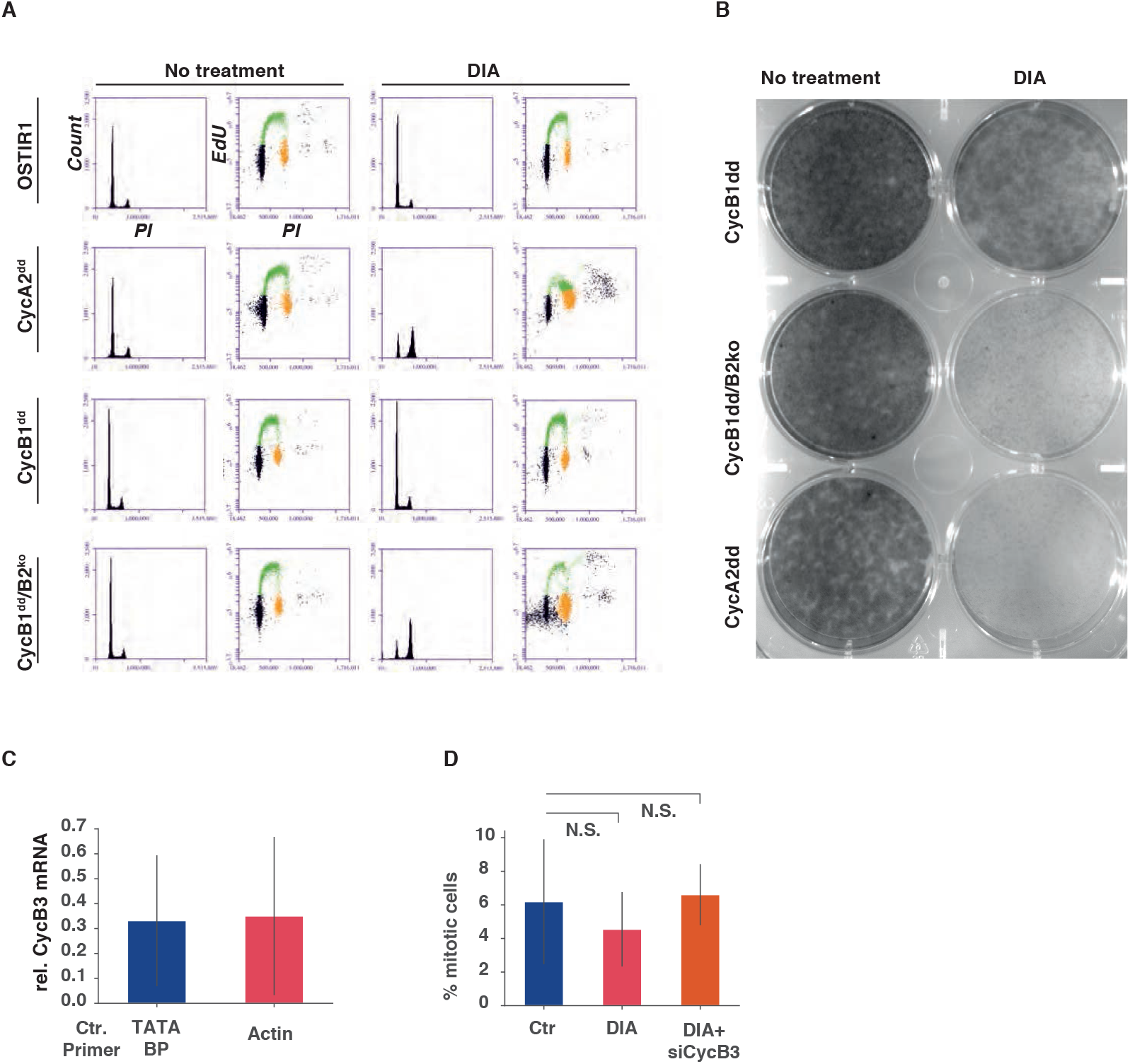
Cell proliferation in the absence of Cyclin A2, and B-type Cyclins. **(A)** Cell cycle analysis 24 hours after DIA treatment. Cells were analysed by EdU labelling, PI staining and FACS analysis. The histograms show the PI intensities while the dot plots show EdU incorporation (Y-axis) vs PI intensity (X-axis). Quantifications of these representative data involving three independent experiments are shown in Figure 1C. **(B)** Cell proliferation of A2^dd^, CycB1^dd^ and B1^dd^/B2^ko^ following mock or DIA treatment. 1000 cells were plated in 6 well plates and incubated for 10 days before methanol fixation and Crytsal Violet staining. **(C)** qPCR analysis of Cyclin B3 mRNA levels, following 72 hours depletion in B1^dd^/B2^ko^ cells. For quantificatioon we used primers to amplify two control mRNAs, TATA binding protein, and Actin. The plot shows the levels of CycB3 siRNA depleted mRNA relative to Ctr siRNA transfected cells. **(D)** Mitotic index measurements of B1^dd^/B2^ko^ cells with the indicated treatemnts. Cells were transfected with siRNA for 72 hours blocked for 24 hours with Thymidine fixed 12 hours after release from Thymidine. Protame and Apcin were added for the final two hours before fixation. Mitotic cells were scored based DAPI staining and on condensed chromosome formation, The bar-plots show mean values of three biological releats (n=100 per repast and sample), Error bars indicate standard deviation.

**Supplementary Figure S4.**
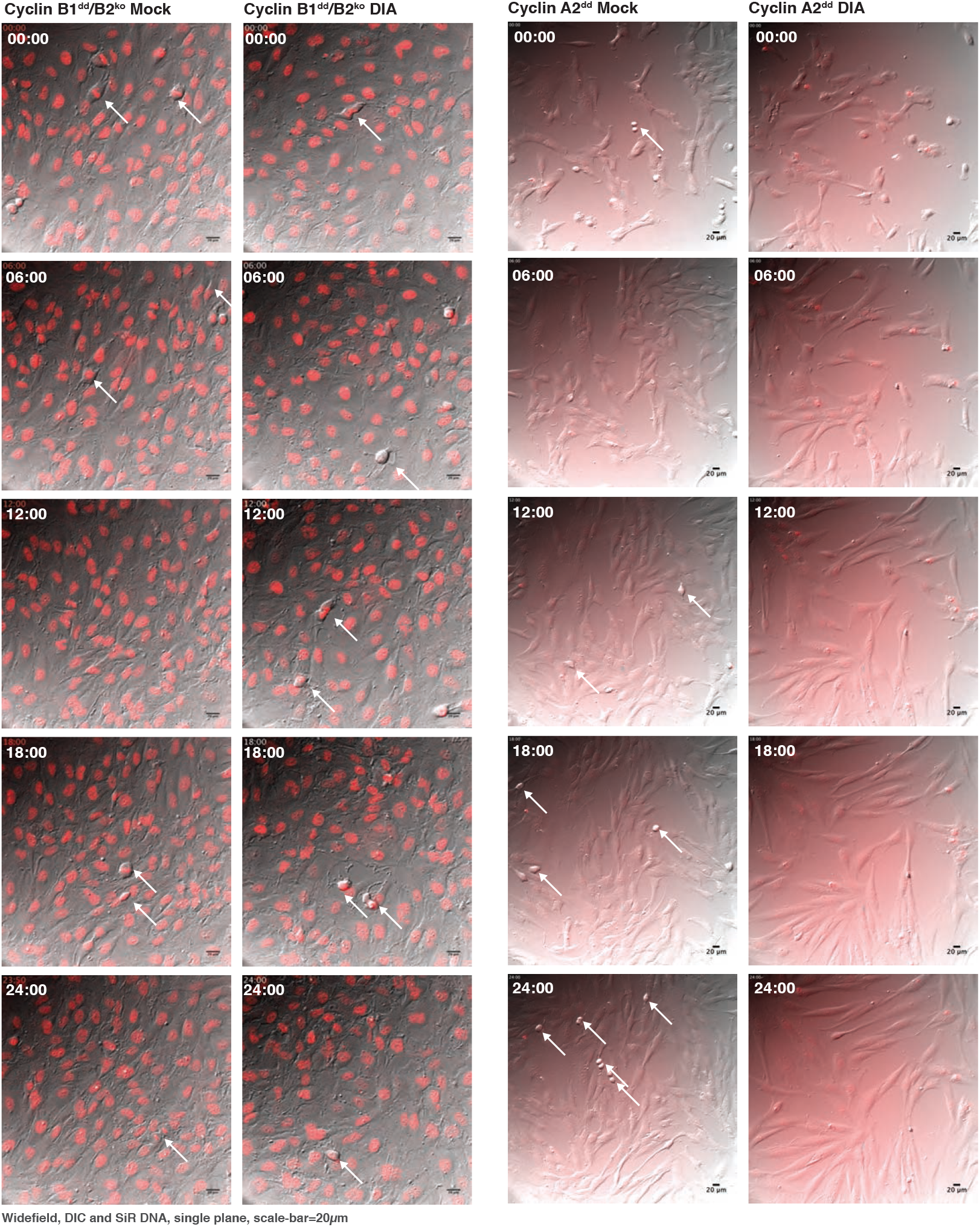
Mitotic progression in RPE-1 cells lacking Cyclins B1 and B2. Differential interference contrast and fluorescence widefield video microscopy of mock or DIA treated SiR-DNA labelled cells with indicated genotypes. Images show a progression of 6 hour time points (time inicated as hh:min).

**Figure S5:**
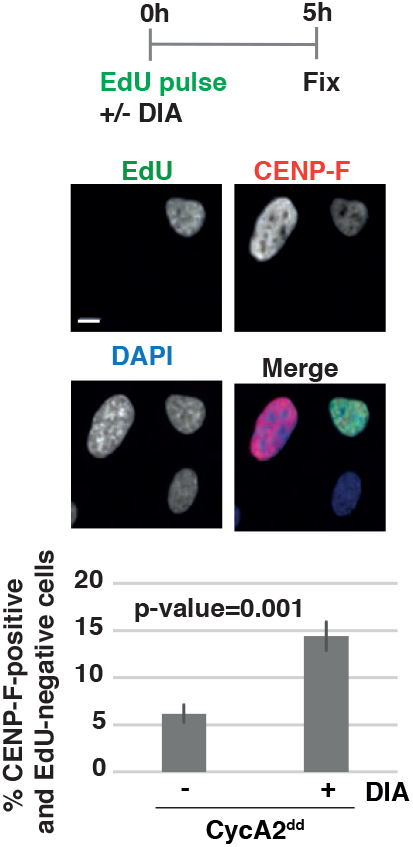
Loss of Cyclin A2 in G2 cells causes block in mitotic progression. A2^dd^ cells were pulsed with EdU to label S-phase then protein degradation was induced by DIA addition for 5h and before fixation and Edu/CENPF/DAPI staining. Cells that were in G2 at the time of DIA or mock treatment were identified as CENPF positive, EdU negative. Mitotic cells were excluded based on nuclei morphology. The bar plots show mean percentage of EdU negative/CENPF positive cells of three repeats, error bars indicate standard deviation. Representative immuno-fluorescence images are displayed (scale-bar 10*μ*m).

**Figure S6.**
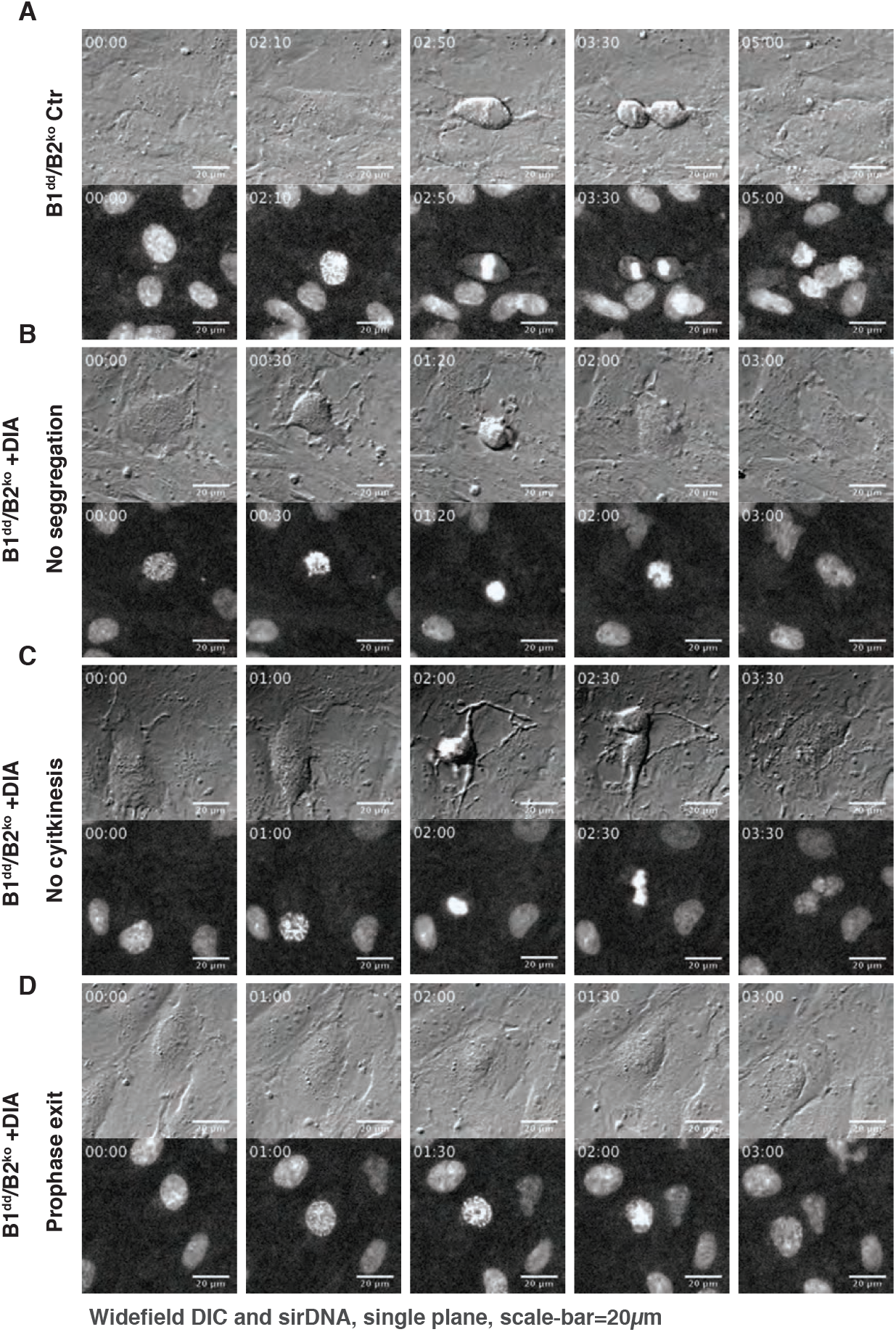
Examples of mitotic phenotype in cells lacking Cyclin B1 and B2 (relating to Figure 3A and B) Stills from live cell imaging of asynchronously dividing B1^dd^B2^ko^ cells (DIC (top) and sirDNA (bottom), time is indicated as hh:min). **(A)** shows a successful cell division in untreated B1^dd^B2^ko^ cells. **(B)** shows DIA treated B1^dd^B2^ko^ cells that fail to segregate sister chromatids and initiate cell division. **(C)** shows DIA treated B1^dd^B2^ko^ cells that attempt segregation and cytokinesis but fail to separate resulting in a binuclear cells. **(D)** shows DIA treated B1^dd^B2^ko^ cells that fails to undergo NEBD and exits mitosis from prophase.

**Figure S7.**
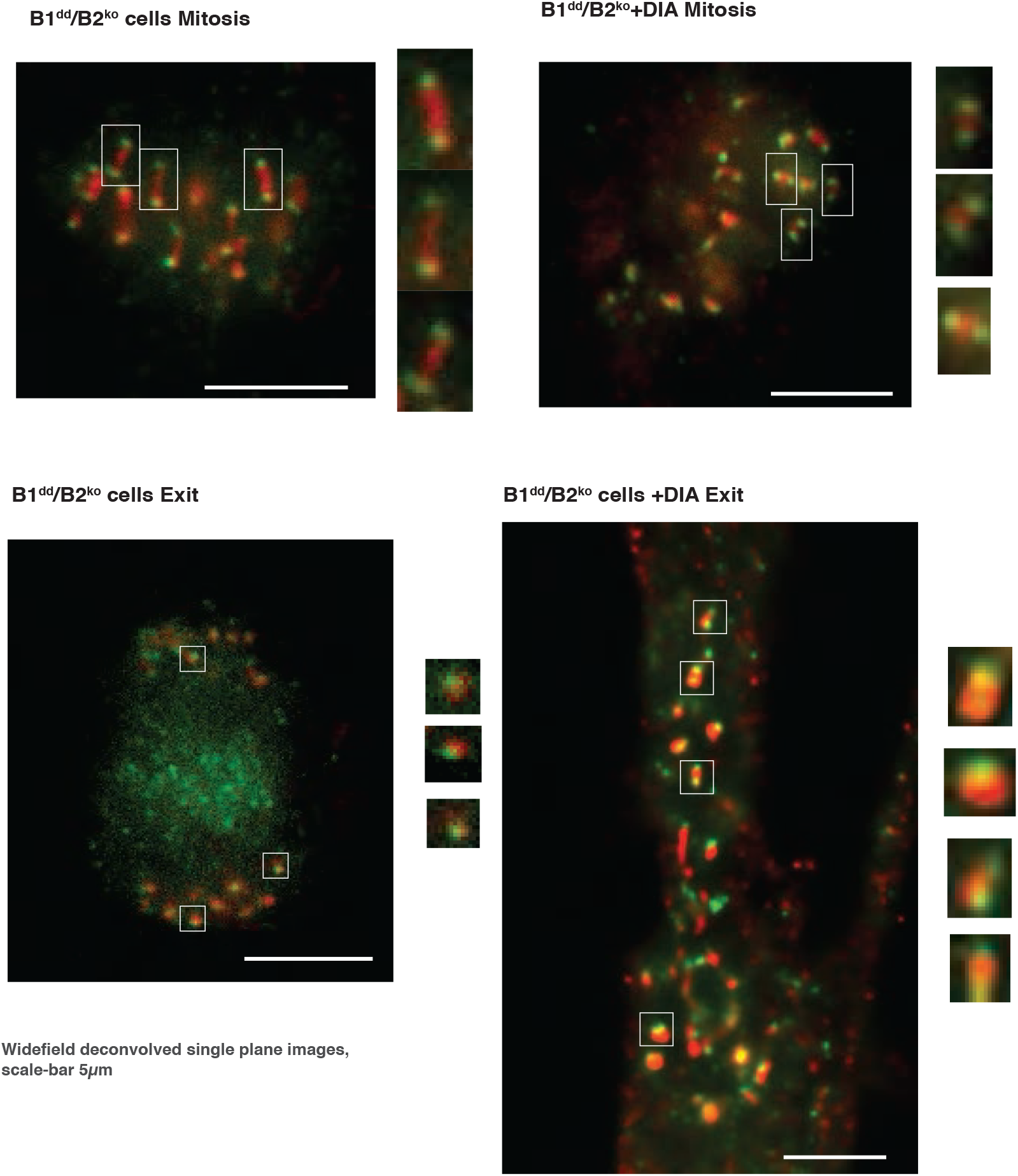
Analysis of Kinetochore tension and segregation. Immuno-fluorescence images from mitotic (Mitosis, top-two panels) and ana/telophase (Exit, bottom two panels) Ctr and DIA treated B1^dd^/B2^ko^ cells stained with anti-CenpA (green) and anti-CenpB (red)-antibodies. For DIA treated cells we determined the exit state by the onset of a cytokinesis furrow, since sister chromatid segregation did not occur. Areas marked by squares are enlarged twofold at the right of each panel.

**Figure S8.**
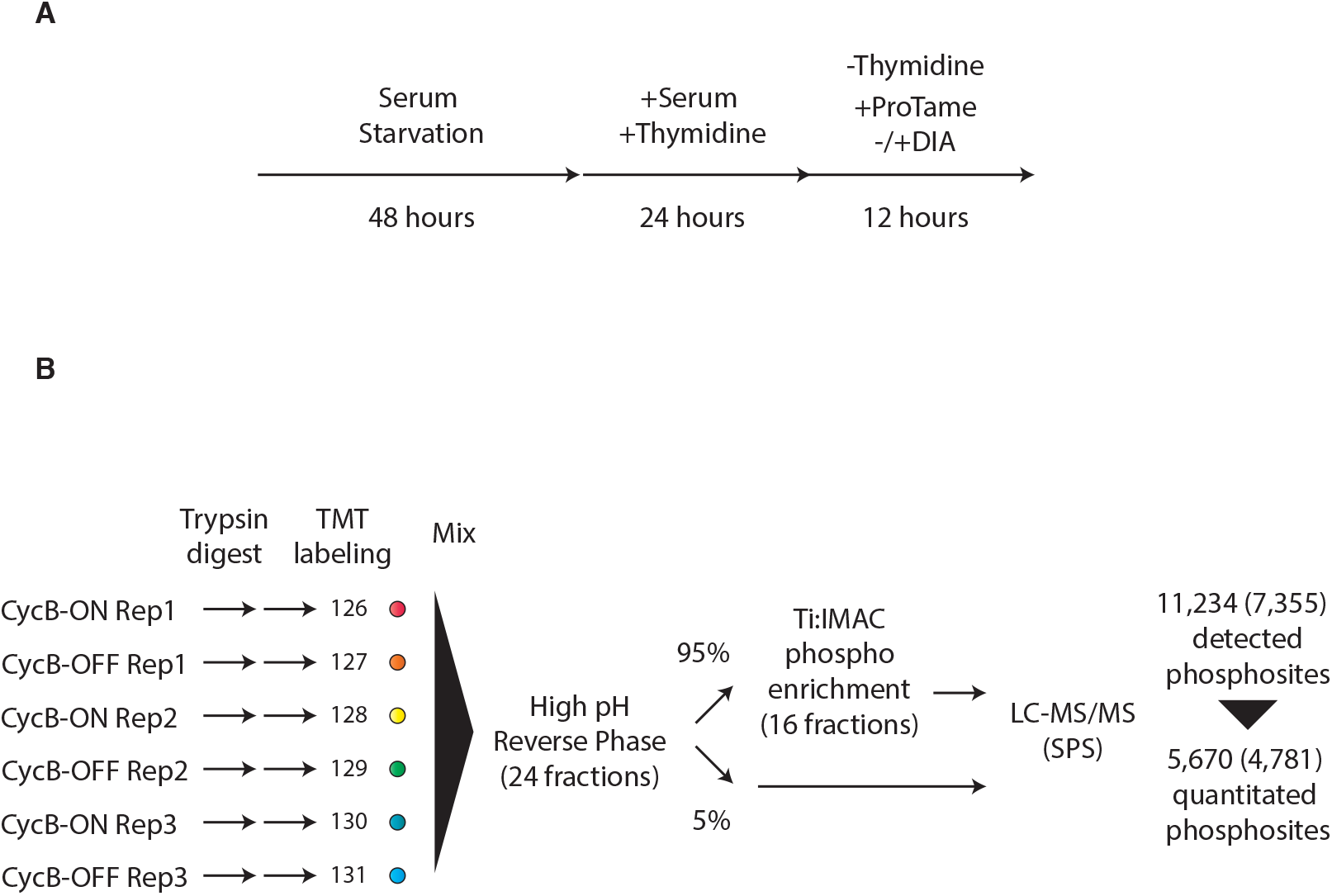
Experimental scheme for phosphorylation-proteomic analysis. **(A)** Cell synchronisation scheme: To obtain an optimal enrichment in mitotic cells we presynchronised B1^dd^/B2^ko^ cells by serum starvation for 48 hours followed by a release in thymidine for 24 hours. Cells were then released into S-phase in the presence or absence of DIA and blocked in mitosis by the addition of ProTame. Mitotic cells were collected by shake-off 12 hours following release. **(B)** Phosphoproteomics workflow. Protein extracts were prepared from the indicated cells, digested with trypsin and labelled with 6-plex TMT reagents. The TMT-labelled peptides were then mixed and subjected to fractionation by high pH reverse phase into 24 fractions. From each fraction, 95% was taken for phosphoenrichment and 5% retained for analysis of unmodified peptides. For phosphoenrichment, the 24 fractions were combined into a total of 16 fractions. Samples (unmodified and phosphoenriched) were then analysed by LC-MS/MS on an Orbitrap Fusion instrument using synchronous precursor selection (SPS). This analysis resulted in detection of 11,234 phosphorylation sites, of which 5,670 were quantitated. The number of class I sites are indicated in brackets (site localisation probability > 0.75).”

**Figure S9.**
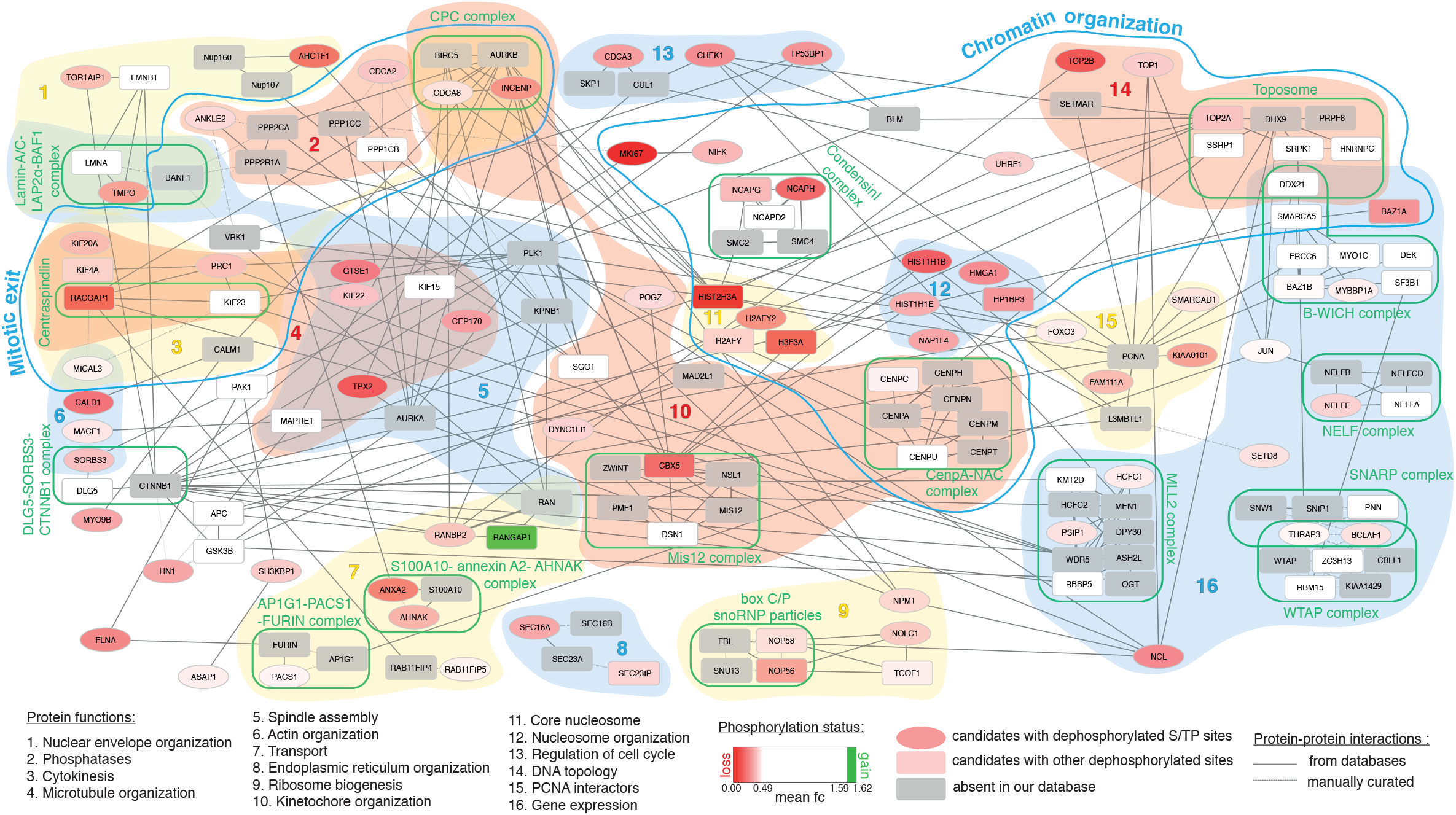
Network analysis of identified of Cyclin B substrates

**Figure S10.**
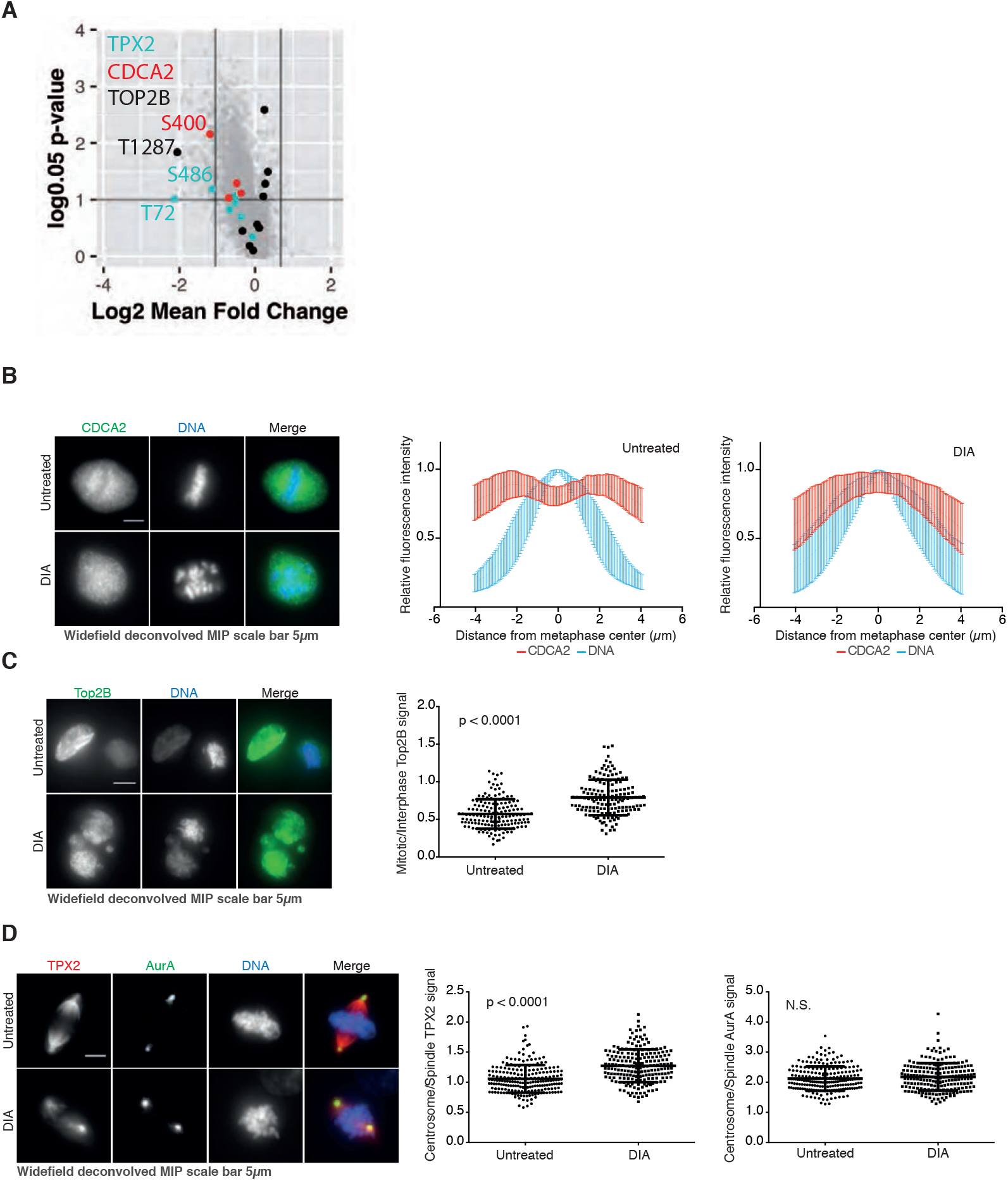
Validation of hits identified in proteomic screen. **(A)** Identified phospho-sites in candidates chosen for validation of the proteomic data-set: Tpx2, CDCA2 (Repoman) and Top2B **(B)** CDCA2 is excluded from mitotic chromosomes in control cells while this exclusion is lost in DIA treated B1^dd^/B2^ko^ cells. The left panel shows immuno-fluorescence images, for quantification we analysed fluorescentce intensity across chromosomes. The data show the mean of 150 cells (from three repeats n=50) and errorbars indicate stdv. **(C)** Top2B levels in mitotic cells are increased in DIA treated B1^dd^/B2^ko^ cells. The left panel shows immuno-fluorescence images, for quantification we segmented mitotic DAPI staining and measured mean intensity of Top2B staining. Three repeats n>50 cells per repeat. **(D)** Tpx2 shows reduced spread on mitotic chromosomes in DIA treated B1^dd^/B2^ko^ cells. The left panel shows immuno-fluorescence images, for quantification we measured mean intensity of Tpx2 staining on the centrosome and spindle and plotted the ratio of these values per cell. Three repeats n>50 cells per repeat. As a control we analysed the centrosome/spindle distribution of AurA that does not change following DIA treatment.

**Figure S11.**
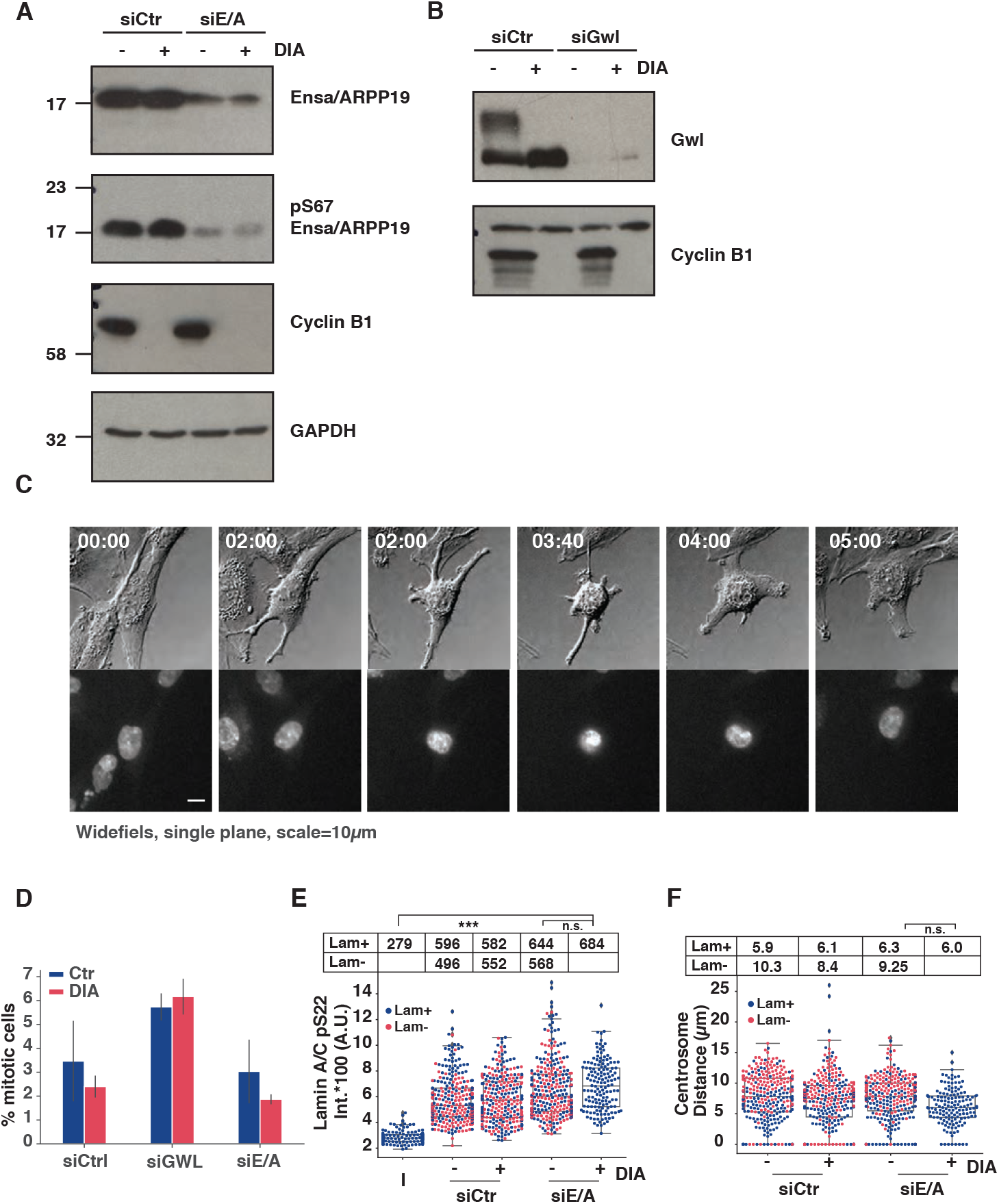
Co-depletion of Ensa/ARPP19 or Greatwall and Cyclin B causes a prolonged prophase. **(A)** Immuno-blot of B1^dd^/B2^ko^ cells transfected with indicated Ctr or Ensa/ARPP19 siRNAs. Cells were blocked in Thymidine 12 hours after siRNA tranfection for 24 hours and released for 10 hours before sample preparation. Ensa/ARPP19 depletion is incomplete under these condition but causes a substantial decrease in pS67 phosphorylation indicating that PP2A:B55 is not inhibited. **(B)** As in (A) with Greatwall siRNA tranfetced cells. Note that DIA treatment causes a substantial change in Gwl phosphorylation but does not affect Ensa/ARPP19 S67 phosphorylation (see Panel A) indicating that Gwl is active in DIA treated cells, but that Cyclin B is required for Gwl phosphorylation at residues not required for activity. **(C)** Widefield Imaging. DIC (Grey), SiR-DNA (b/w), of Ensa/ARPP19 siRNA transfected DIA treated B1^dd^/B2^ko^ cells. Time is indicated in hh:min. **(D)** Quantification of mitotic index of siRNA transfected P/A synchronised cells (See Figure 4B for representative images. **(E)** Intensity of anti-Lamin A/C pS22 antibody staining in siRNA transfected and DIA treated B1^dd^/B2^ko^ cells. The swarm blots classify data from mitotic cells with intact (blue) or disassembled (red) Lamin A/C. The table above the panel indicates the median values and selected p-values are indicated **(F)** Same as in (E) for centrosome distance in mitotic cells (Data for (E and F) are from three repeats, n>50 per repeat).

**Figure S12.**
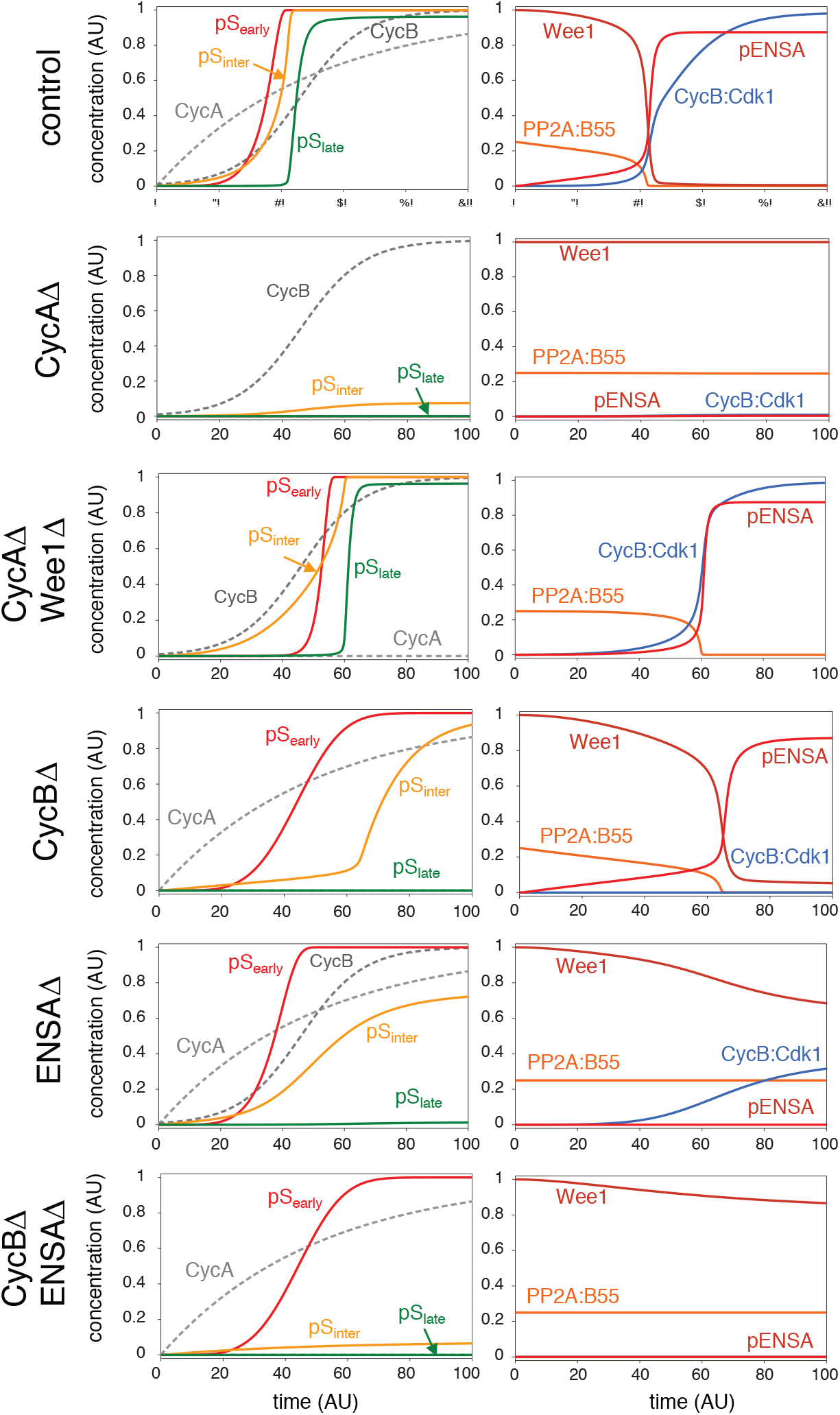
Numerical Simulation of mitotic substrate phosphorylation network in response to rising Cyclin levels. The model simulates mitotic entry experiments where cyclin levels become stabilized by APC/C inhibition. Each row corresponds to one of the experimental conditions used in this work (control, CycA depletion etc.). Left columns: Cyclin levels and phosphorylated mitotic substrates (early, intermediate and late). Right columns: temporal changes of some mitotic regulators. Changes of Cdc25P and GwlP (not shown) are similar to phosphorylated ENSA (pENSA).

## Supplementary Material

### A) Supplementary Videos

**Supplementary Video 1**

B1^dd^/B2^ko^ cells expressing FusionRed-Histone H2B (green) and Mis12-GFP (white) and labelled with SiR-Tubulin (red) going through mitosis. Time is indicated as hh:min. Scale bar represents 5μm. Images are airy scan reconstructions and maximum intensity projections of seven stacks taken at 2μm intervals.

**Supplementary Video 2**

DIA treated B1^dd^/B2^ko^ cells expressing FusionRed-Histone H2B (green) and Mis12-GFP (white) and labelled with SiR-Tubulin (red) going through mitosis. Time is indicated as hh:min. Scale bar represents 5μm. Images are airy scan reconstructions and maximum intensity projections of seven stacks taken at 2μm intervals.

**Supplementary Video 3**

B1^dd^/B2^ko^ cells expressing FusionRed-Histone H2B (green) and Aurora-B-GFP (white) and labelled with SiR-Tubulin (red) going through mitosis. Time is indicated as hh:min. Scale bar represents 5μm. Images are airy scan reconstructions and maximum intensity projections of seven stacks taken at 2μm intervals.

**Supplementary Video 4**

DIA treated B1^dd^/B2^ko^ cells expressing FusionRed-Histone H2B (green) and Aurora-B-GFP (white) and labelled with SiR-Tubulin (red) going through mitosis. Time is indicated as hh:min. Scale bar represents 5μm. Images are airy scan reconstructions and maximum intensity projections of seven stacks taken at 2μm intervals.

**Supplementary Video 5**

Mitotic progression in B1^dd^/B2^ko^ cells transfected with Ctr siRNA with or without DIA treatment and Ensa/ARPP19 siRNA transfected B1^dd^/B2^ko^ cells without DIA treatment. 12 hours after siRNA transfection cells were blocked for 24 hours by Thymidine, released from the arrest and imaged 10 hours after release. The cells are expressing FusionRed-Histone H2B (red) and are labelled with SiR-Tubulin (white). Time is indicated as hh:min. Scale bar represents 5μm. Images are airy scan reconstructions and maximum intensity projections of seven stacks taken at 2μm intervals.

**Supplementary Video 6**

Prophase arrest and slippage in B1^dd^/B2^ko^ cells transfected with Ensa/ARPP19 siRNA treated with DIA following release from Thymidine arrest. The cells are expressing FusionRed-Histone H2B (red) and are labelled with SiR-Tubulin (white). Time is indicated as hh:min. Scale bar represents 5μm. Images are airy scan reconstructions and maximum intensity projections of seven stacks taken at 2μm intervals.

### B) Supplementary Tables

**Supplementary Table 1**

All detected phosphorylation sites. Columns include protein identifiers, the site localisation probabilities, positions in protein, positions in peptide sequence, the identification score, the posterior error probability (PEP), the localisation score, the normalised TMT reporter intensities for the phosphorylated peptides, the normalised TMT reporter intensities for the unmodified protein, ratio of phosphorylated/unmodified, the mean fold change (3 biological replicates), the p-value (t-test, uncorrected), and columns indicating whether several linear motifs are found in the phosphorylated peptide sequence (e.g. CDK consensus motif, etc.)

**Supplementary Table 2**

Significantly changing phosphorylation sites as determined by 2x-fold change and 0.05 p-value cutoffs. Columns include protein identifiers, the site localisation probabilities, positions in protein, positions in peptide sequence, the identification score, the posterior error probability (PEP), the localisation score, the normalised TMT reporter intensities for the phosphorylated peptides, the normalised TMT reporter intensities for the unmodified protein, ratio of phosphorylated/unmodified, the mean fold change (3 biological replicates), the p-value (t-test, uncorrected), and columns indicating whether several linear motifs are found in the phosphorylated peptide sequence (e.g. CDK consensus motif, etc.)

### C) Material and Methods

#### Tissue Culture and Chemical Reagents

hTERT RPE-1 cells were obtained from ATCC (cat. CRL-4000) and were grown at 37°C with 5% CO2 in DMEM/F12 (Sigma-Aldrich) media containing 10% fetal bovine serum (FCS) and 1% penicillin–streptomycin.

> Drugs used for this study and working concentrations were:
>
> PD0166285 (0.5μM) (Stratech Scientific Limited, S8148-SEL)
>
> Apcin (26μM) (Sigma-Aldrich SML1503)
>
> ProTame (6μM) (Bio-techne I-440-01M)
>
> Asunaprevir (3μM) (BMS-650032, Bioquote, A3195)
>
> indole-3-acetic acid (IAA, 500μM) (Sigma Aldrich I5148)
>
> Doxycyclin (1μg/ml) (Takara-bio, Clontech 631311)
>
> Thymidine (4mM) (Sigma Aldrich T1895)
>
> SiR-DNA (50-100nM) (Tebu-bio Ltd. SC007)
>
> SiR-Tubulin (50-100nM) (Tebu-bio Ltd. SC002)

#### Antibodies

| Antigen | Host | Company | Cat. Number | secondary ABs | Company | Cat. Number |
| --- | --- | --- | --- | --- | --- | --- |
| Alpha-tubulin | mouse | Abcam | nr. 7291 | Alexa647 donkey anti-rabbit | Invitrogen | A31573 |
| Gamma-tubulin | rabbit | Abcam | ab84355 | Alexa555 donkey anti-goat | Invitrogen | A21432 |
| Pericentrin | rabbit | Abcam | ab4448 | Alexa488 donkey anti-mouse | Invitrogen | A21202 |
| CREST | human | ImmunoVision | HCT-0100 | HRP Goat anti-mouse | DAKO | P0447 |
| Lamin A/C | goat | Santa Cruz | sc-6215 Lot l2111 | HRP Goat anti-Rabbit | Bethyl | A120-201P |
| pLamin A/C S22 | rabbit | Cell Signaling Technology | 2026S |  |  |  |
| pCdk1-substrates | rabbit | Cell Signaling Technology | 9477S |  |  |  |
| Aurora B | rabbit | Abcam | ab2254 |  |  |  |
| Kl67 | rabbit | Abcam | ab16667 |  |  |  |
| Cyclin A2 | mouse | Abcam | ab38 |  |  |  |
| Cyclin B1 | mouse | Abcam | ab72 |  |  |  |
| Cyclin B2 | mouse | SC | sc-28303 |  |  |  |
| Cyclin B3 | mouse | SC | sc-515887 |  |  |  |
| Cyclin B3 | rabbit | ThermoFisher | PA5-37254 |  |  |  |
| myc | mouse | Abcam | ab32 |  |  |  |
| GAPDH | mouse | Genetex | GTX627408 |  |  |  |
| Aurora A | mouse | Abcam | ab13824 |  |  |  |
| Aurora B | rabbit | Abcam | ab2254 |  |  |  |
| ENSA | rabbit | Abcam | ab180513 |  |  |  |
| pENSA (ser67) | rabbit | CST | nr. 5240 |  |  |  |
| Greatwall | rabbit | Sigma | HPA02717 |  |  |  |
| Tpx2 | rabbit | Bethyl | A300-429A |  |  |  |
| Repoman (CDCA2) | rabbit | Sigma | SAB4500599 |  |  |  |
| CenpA | mouse | Abcam | ab13939 |  |  |  |
| CenpB | rabbit | Abcam | ab25734 |  |  |  |
| Topo2B | mouse | BD Transd. Labs | nr. 611492 |  |  |  |

#### Generation of endogenously tagged cell lines

gRNAs were designed using Benchling CRISPR tool (https://benchling.com/). The sequences 5’ ACTAGTTCAAGATTTAGCCA 3’, 5’ TGTTTCTAAAACCATCAAGT 3’, 5’ GCACACTCACCGTCGGGCGT 3’ and 5’ GACCTGCTACAGGCACTCGT 3’ were chosen for cyclin B1, cyclin A2, cyclin B2 and Rosa26 respectively, and introduced into the vector pSpCas9(BB)-2A-Puro (PX459) V2.0 following the protocol described in (*1*). PX459 acquired from Feng Zhang via Addgene (plasmid # 48139). Indel mutations in Cyclin B2 were confirmed by Sanger sequencing as two frameshift mutations downstream of the initiating ATG in the CCNB2 gene (CTCGACGCCCGACG-GTGAG and CTCGACGCC-C-GACGGTGAG with the missing residues marked by hyphenation). The Puromycin resistance in hTERT RPE-1/OSTIR1 cells was removed using CRISPR using the following gRNA sequence: 5’ AGGGTAGTCGGCGAACGCGG 3’. To make the targeting template, Gibson assembly was used to assemble into NotI-digested pAAV-CMV vector (gift from Stephan Geley, University of Innsbruck, Austria) the fragments in the following order: the left arm, a linker (5’ CGCCTCAGCGGCATCAGCTGCAGGAGCTGGAGGTGCATCTGGCTCAGCGGCAGG 3’), mAID 3, SMASh 5, T2A-neomycin and the right arm. To get CRISPR-resistant constructs, the following sequences were mutated as followed: ACTAGTTCAAGATTTAGCCAAGG by AtTAGTcCAgGAccTAGCtAAaG for cyclin B1 and CCATCAAGTCGGTCAGACAGAAA by CCATgAtGaCGcTCAcACAGttA for cyclin A2. Mutations (lowercase letters) are silent and preferential codon usage was taken into account. For inducible expression of OsTIR1 we used the construct described in (*2*) combined it with a bleomycin/zeocin resistance marker and cloned it into a Rosa26 targeting construct. Integration was confirmed by genomic PCR (Supplementary Fig. 1b). To generate stable clones, 106 hTERT immortalised RPE-1 cells were transfected with 0.5μg of gRNA/Cas9 expression plasmid and 1.5μg of targeting template using Neon transfection system (Invitrogen), with the following settings: 10μL needle, 1350V, 20ms and 2 pulses. Clones were incubated for 3 weeks in media containing 1mg/mL of neomycin (Sigma-Aldrich), 5μg/mL blasticidin (Gibco) or 500μg/mL zeocin (Invivogen) and selected clones were screened by western blot.

#### Generation of PCNA tagged cell lines

AAV-293T cells (Clontech) were seeded into a T75 flask one day before transfection, such that they were 70% confluent on the day of transfection. Cells were transfected with 3μg each of pAAV-mRuby-PCNA (*3*) pRC and pHelper plasmids, and 20μl Lipofectamine 2000, diluted in 3ml OPTI-MEM (Gibco). Lipofectamine/DNA mixture was added to cells in 7ml of complete medium (DMEM with 10% FBS and 1% penicillin-streptomycin). Cells were incubated at 37’C for 6 hours and before medium was replaced with 10ml of complete medium. Three days post transfection, medium and cells were transferred to a 50 ml falcon tube. Cells were lysed with three rounds of freeze-thaw. The sample was centrifuged at 10,000 x g for 30min at 4’C. Supernatant containing AAV particles was collected and either used immediately or aliquoted and stored at −80’C. Cyclin A2^dd^ cells were plated one day before transduction, such that they were 40% confluent for transduction. Cells were washed twice in PBS and incubated in 5ml of complete medium plus 5ml of AAV-mRuby-PCNA containing supernatant for 48 hr. Cells were expanded for a further 48 hr followed by FACS sorting using a BD FACSMelody sorter according to the manufacturer’s instruction.

#### Generation of cell lines stably expressing fluorescent protein markers

For rapid generation of multiple fluorescent protein tagged cellular markers we cloned a sequence of P2A-ScaI-mEmeraldT2A-Balsticidin resistance marker into the pFusionRed-H2B expression construct (Evrogen, FP421). the ScaI site was then used to clone Mis12 and AurB in-frame with the preceding P2A and the following T2A sequence. Cyclin A2dd and B1ddB2ko cells were transfected with 2μg of the expression plasmids by NEON electroporation (Invitrogen) and grown for two weeks in medium containing 5μg/mL blasticidin (Gibco). Fluorescent protein expressing cell lines were isolated by FACS sorting using a BD FACSMelody sorter according to the manufacturers instruction.

#### Genomic PCR

Genomic DNA was extracted using DNeasy Blood and Tissue Kit (Qiagen) according to the instructor’s recommendation then DNA was amplified with Phusion High Fidelity DNA polymerase (New England Biolabs) using the following primer pairs to check genomic integration: 5’ CTGCATTCTAGTTGTGGTTTGTCCA 3’ and 5’ ACTATGACCCACGCAGTACAA 3’ (TetON-OsTIR1 into Rosa26 locus), 5’ ATTGCTGAAGAGCTTGGCGG 3’ and 5’ TCACACCATTCAAGCACCTGTA 3’ (degron tags into Cyclin A2 locus), 5’ CTGAGCGGAAAACCTGCTATC 3’ and 5’ CTGAACGAACAGGGGAAATGGTT 3’ (degron tags into Cyclin B1 locus), 5’ TGGTGGAA-GATTGGTGGCTC 3’ and 5’ CTGCTTCTGGCATGGCTTTC 3’ (internal amplification control in Kif23 locus). Samples were loaded onto 1% agarose gel (Fisher) in 1X TAE buffer and containing ethidium bromide then imaged using Ingenius, Syngene Bio-imaging apparatus.

#### RT-qPCR

cDNA was prepared from extracted mRNA (RNeasy kit, Qiagen) using oligo d(T) (Ambion), murine RNase inhibitor (New England Biolabs) and M-MuLV reverse transcriptase (New England Biolabs) following the manufacturer’s directions. Samples were then treated with RNase A (Sigma) and cleaned using QIAquick PCR purification kit (Qiagen). Next, qPCR was carried out with HOT FIREPol EvaGreen qPCR Mix Plus (no Rox, Solis BioDyne) and primers for cyclin B3 (5’ AAGACACTGACCTTGTCCCG 3’ and 5’ AGAGGGCCAGGAGTAAGGAG 3’) and the loading controls: actin (5’ GAAGTGTGACGTGGACATCC 3’ and 5’ CTCGTCATACTCCTGCTTGC 3’) and TATA-binding protein (5’ CACGAACCACGGCACTGATT 3’ and 5’ TTTTCTTGCTGCCAGTCTGGAC 3’). The reaction was performed on a PCR Stratagene MX3005P system. Fold change in expression over control was calculated using the delta-delta Ct method (*4*).

#### Proliferation assay

1000 cells per well were seeded and treated or not with 1μg/mL of doxycycline, 3μM of Asv and 500μM of IAA for 1 week then cells were fixed and stained with a solution containing 0.05% crystal violet (Sigma-Aldrich), 1% formaldehyde (Sigma-Aldrich), 1% methanol in PBS for 10min.

#### FACS analysis

Cells were incubated with 10μM of EdU for 1h before being harvested and fixed in 70% ethanol. Next, EdU was labelled with the fluorophore Alexa 647 using Click-iT EdU Imaging Kit (Invitrogen) and DNA were stained with 5μg/mL of propidium iodide (Fluka) and 150μg/mL of RNAseA (Sigma-Aldrich). FACS profiles were obtained from data acquired on Accuri C6 Flow cytometer (BD Biosciences).

#### Cell synchronisation

To enrich mitotic cells and block cells in metaphase, we used a single 24 hour Thymidine block release. Apcin (25μM) and ProTame (6μM) were added 12 hour after the release for 2 hours before fixation and analysis. For more efficient mitotic enrichment to collect cells for proteomic or immunoblotting analysis we pre-synchronised cells by serum starvation for 72 hours, followed by a 24 hour release into serum and Thymidine containing medium, followed by release from thymidine. ProTame (6μM) was added 12 hours after release for two to four hours and mitotic cells were collected by shake off.

#### siRNA transfection

siRNAs were resuspended to 20μM stock concentration. 1.5ml 20×104 cells were reverse transfected with siRNA diluted in 500μL MEM (Gibco) media containing (10μl) siRNA-MAX reagent (Invitrogen) and a final concentration of 80nM siRNA. After 7h-O/N incubation the media was changed to the standard growth media. The cells were subsequently used for analysis after 48h depending, or 72 hours in the case of CCNB3. The following siRNAs were used for this study siGWL (Qiagen Ltd SI02653182 *HsMASTL7* FlexiTube), siARPP19 (Dharmacon L-015338-00), siENSA (Dharmacon L-01182-00), siCCNB3 (Dharmacon L-003208-00-0005), All Stars negative control siRNA (Qiagen 1027280). In the case of Gwl and Ensa/ARPP siRNA transfections cells were synchronised 12 hours after siRNA transfection by a 25 hour single Thymidine block, and analysed 10-12 hours after release.

#### Immuno-blotting

Cells were pre-treated for 2h with 1μg/mL of doxycycline (Clontech) then 3μM of asunaprevir (Asv; ApexBio) and/or 500μM IAA were added to the media for indicated time periods. Cells were harvested and lysed in EBC buffer (50mM Tris pH 7.5, 120mM NaCl, 0.5% NP40, 1mM EDTA, 1mM DTT, Protease and Phosphatase inhibitors (Complete and PhosStop; Roche Diagnostics)) then mixed with 5x sample buffer (0.01% bromophenol blue, 62.5mM Tris-HCl pH 6.8, 7% SDS, 20% sucrose and 5% β-mercaptoethanol). The samples were sonicated then boiled at 95°C for 5min. Samples were analysed by western blotting and the signal was detected using Immobilon Western Chemiluminescent HRP substrate (Millipore). The intensity of cyclins A2 and B1 signals were quantified using ImageJ software. GAPDH was used to normalize the samples.

#### Immuno-fluorescence

Cells were fixed for 10 minutes with 3.7% formaldehyde (Sigma-Aldrich) in PBS, washed in PBS and permeabilised with PBS containing 0.5% NP40 for 10 minutes. Following 30 minutes blocking in 3% Bovine Serum Albumin (BSA, Sigma Alderich) cells were labelled with the indicated antibodies diluted in 3%BSA/PBS. Secondary antibodies were labelled with Alexa Fluor dyes purchased from Invitrogen. Cell nuclei were counterstained with 4,6-diamidino-2-phenylindole dihydrochloride (DAPI; Sigma-Aldrich) then slides were mounted with ProLong Diamond Antifade Mountant (Invitrogen).

#### Cold treatment

Cells were grown on coverslips and were or not pre-treated with 1μg/mL of doxycycline for 2h then 3μM of Asv and 500μM of IAA were added for 4h. Next, the cells were incubated on ice for 10 min. For microtubule regrowth assay, the cells were re-incubated at 37°C for indicated times. Fixation was carried out with PHEM (60mM Pipes, 25mM HepesKOH pH 7.0, 5mM EGTA, 4mM MgSO4, 0.5% NP40 and 3.7% formaldehyde) for 5 min followed by 95% ice-cold methanol/5mM EGTA for 5 min. The samples were then subjected to immuno-fluorescence, widefield microscopy an deconvolution.

#### Chromosome spreads

Cells were synchronised by single Thymidine release and Protame/Apcin arrest as described above. Mitotic cells were collected by shake off and resuspended in warm 75mM KCl solution for 10min before centrifugation. After aspirating KCl, Methanol:Acetic acid (3:1) was added to cells and incubated at RT for 5min. Cells were centrifuged and resuspended in 300-500l remaining solution and dropped on glass coverslips. They were allowed to dry for 1h prior to additional 4% Formaldehyde fixation for 10min followed by immuno-fluorescence using Ki67 antibodies.

#### Widefield live cell microscopy

Cells were seeded on μ-slide from Ibidi. Cells were pre-treated with 2μg/mL of doxycycline for 2h before imaging and 3μM of Asv and 500μM of IAA were added at the beginning of the imaging. To observe the nucleus, cells were pre-treated with 50nM SiR-DNA (Spirochrome) for 2h. To analyse mitotic entry, time-lapse microscopy was performed in an environmental chamber (Digital Pixels, Microscopy Systems & Solutions) heated at 37°C with 5% CO2 supply using an Olympus IX71 equipped with Orca-flash4.0LT camera and a LUCPlanFLN, NA 0.45, 20X objective, or 0.64 NA, LUCPlanFLN 40x lens and 2 × 2 binning. Cells were imaged using differential interference contrast (DIC), and fluorescent illumination using a Lumencor Spectra LED light source, and 640/40 Excitation, 705/72 Emission filters. Images were acquired every 5min using Micro-Manager v1.4 software and cells that rounded-up and condensed their nucleus were scored.

#### Widefield immuno-fluorescence microscopy and deconvolution

Slides were imaged using an Olympus IX70 equipped with CoolSNAP HQ2 camera and an UApo N 340 NA 1.35, 40X oil immersion objective controlled by Micro-Manager v1.4 software. Stacks were taken at 0.2μm intervals and deconvolved using the Huygens Classic Maximum Likelihood Estimation algorithm (Scientific Volume Imaging) based on a measured point spread function.

#### Airyscan confocal microscopy (Zeiss)

Confocal imaging was performed on the Airy scan module of Zeiss LSM880 (Carl Zeiss AG^®^, Germany) with Plan-Apochromat 63x/1.4 Oil objective in a live support chamber (Digital Pixels, Microscopy Systems & Solutions) at 5%CO2 and 37°C. The ‘Fast’ scanning mode was used to speed up the imaging. At each imaging position, 7 axial slices with 2um interval were taken in three fluorescence channels including EGFP, RFP and Cy5. The excitation wavelengths were 488nm, 561nm and 633nm. The emission filters in front of the Airyscan detectors were BP495-550, BP495-620 and LP645 respectively. Each image was accumulated from four scans to minimize laser induced photo-bleaching and photo-toxicity. The time lapses were set to last over 18 hours with 5 mins interval. Raw images were processed using Airyscan processing algorithm in Zen (Carl Zeiss AG^®^, Germany). The processed image sequence was then maximum projected.

#### Spinning Disc Confocal imaging on Operetta (Perkin Elmer)

High throughput imaging was performed on Operetta CLS (PerkinElmer Ltd@) in confocal mode with 40x/0.9 water objective. Cells were cultured and imaged in PerkinElmer CellCarrier Ultra 96 well plates. At each imaging point, 3 axial slices with 4 μm interval were scanned every 5 mins in three fluorescence channels including EGFP, RFP and Cy5. The imaging lasted around 18 hours in 5% CO2 and 37°C live environment. The image sequence was maximum projected, and then analyzed using Harmony software.

#### Image segmentation and quantification of immuno-fluorescence

Regions of interest were either manually generated in ImageJ or using Harmony segmentation algorithms. Mean and Sum Intensity were estimated in ImageJ or Harmony and plotted using Python Pandas, Matplotlib and Seaborn APIs (https://www.python.org,https://matplotlib.org, https://seaborn.pydata.org)

#### High pH reverse phase chromatography and phospho-enrichment

Control and DIA-treated cells were lysed in 2% SDS in Dulbecco’s PBS containing protease and phosphatase inhibitors (Roche; mini-cOmplete protease inhibitors EDTA-free, PhosStop). Lysates were homogenised by sonication using a probe sonicator (Branson sonifier, 20% power, 30 s). 200 μg protein was reduced with 25 mM TCEP (Thermo Pierce), alkylated with N-ethylmaleimide (Sigma, 55 mM prepared fresh) and then precipitated using the chloroform-methanol method. The protein precipitate was resuspended in 0.1 M triethylammonium bicarbonate (TEAB), pH 8.5 and digested first with 2 μg LysC (Wako) for 4 hours and then 2 μg trypsin (Thermo Pierce) overnight. The digest was then reacted with 6-plex TMT reagents (Thermo Pierce) using the manufacturer’s recommended protocol. The TMT-labelled peptides were then mixed.

Peptides were separated off-line by high pH reverse phase chromatography using an Ultimate 3000 high performance/pressure liquid chromatography system (HPLC) (Thermo Dionex) equipped with a BEH 4.6 cm x 150 mm BEH C18 column (Waters) using the following mobile phases: A) 10 mM ammonium formate, pH 9.3 and B) 10 mM ammonium formate, pH 9.3 in 80% acetonitrile (ACN). Peptides were eluted using a linear gradient into 24 fractions. 5% of each fraction was dried and retained aside for total proteome analysis. The remaining 95% was taken forward for phosphoenrichment.

Phospho-enrichment was performed using magnetic Ti:IMAC beads according to manufacturer’s instructions (Resyn Biosciences), similar to previously described (*5*). Dried fractions were resuspended in load buffer (80% ACN, 5% TFA, 5% glycolic acid) and combined into 16 fractions in the following manner: F17+F1, F18+F10, F19+F11, F20+F12, F21+F14, F23+F15, F24+F16. Enriched phosphopeptides were concentrated by evaporation.

#### LC-MS/MS and MS data analysis

Peptides were resuspended in 5% formic acid and separated by reverse phase chromatography on a Ultimate3000 RSLCnano HPLC (Thermo Dionex) equipped with an EasySpray 75 μm x 50 cm column (Thermo Scientific). Peptides were eluted using a linear elution gradient from 2 to 35% B over 2 hours using the following mobile phases: A) 0.1% formic acid, and B) 80% ACN + 0.1% formic acid. Peptides were then ionised and desolvated in an EasySpray electrospray source with a 2.5 kV voltage applied between the emitter and the ion transfer capillary front-end of a Fusion Linear Ion Trap-Orbitrap mass spectrometer (Thermo Scientific). The acquisitions were data-dependent. For MS1 scans, ions from 350 to 1400 m/z were selected using the quadrupole and m/z scanned using the Orbitrap at 120,000 resolution. For MS2 scans, the instrument was operated in top speed mode. Precursors were selected using a 1.6 m/z window for CID fragmentation at 30 normalised collision energy units followed by m/z measurement in the linear ion trap. The top 3 MS2 fragments were then selected for synchronous precursor selection (SPS) using an isolation window of 2 m/z and a maximum injection time of 300 ms. The MS2 fragments were then subjected to further HCD fragmentation at 55 normalised collision energy units. An MS3 scan of the HCD fragments was then performed in the Orbitrap at 60,000 resolution using a scan range of 100 to 150 m/z.

Peptide identification and quantitation was performed using MaxQuant (version 1.5.6.5) (*6*), which incorporates the Andromeda search engine (*7*). Default parameters for TMT quantitation were used in MaxQuant, allowing for the following variable modifications: Phospho(STY), protein N-terminal acetylation, glutamine to pyro(glutamate) conversion, and deamidation of Gln and Asn. The human reference proteome from UniProt (accessed on July 25, 2016) was used as the sequence database for the identification. Raw Data are available via ProteomeXchange with identifier PXD012100

Post-MaxQuant processing was performed using R (version 3.5.0). Potential contaminant proteins and hits to the reverse database were removed from consideration. TMT reporter were normalised by the median protein TMT reporter intensities to correct for unequal loading and also corrected for isotope contamination. Phosphorylation site ratios were then calculated for each of the three biological replicates and normalised to the ratios measured for the total protein. Fold-change cutoffs at 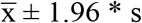 were determined by modelling log ratios as a normal distribution. Motif analysis was performed by mapping regular expressions downloaded for phosphorylation consensus motifs from ELM.

#### Functional classification of the candidates

Outliers were analysed for biological process ontology. Annotations were extracted from the Gene Ontology database using QuickGO. Only qualifier ECO:0000269 experimental evidence used in manual assertion was considered. Biological process ontology verified by direct assay or mutant phenotype from Uniprot database was used to complement the protein classification.

#### Protein interaction map

The interactome was built from human protein-protein interaction (PPI) databases. All Outliers were considered to build a PPI map using the software Cytoscape 3.5.1. Human PPIs from databases were retrieved using the plugin Bisogenet. Small-scale studies were considered and in vivo interactions or PPIs from direct complexes were displayed. The PPI map was complemented using data concerning protein complexes obtained from Uniprot and Corum (http://mips.helmholtz-muenchen.de/corum/). Only PPIs relevant or possibly related to mitosis and proteins present in several protein complexes were shown.

#### Mathematical modelling of mitotic substrate phosphorylation

We have extended our previous model (*8*) with three different (early, intermediate and late) mitotic substrates showing different sensitivities to Cyclin:Cdk1 complexes and PP2A:B55 phosphatase (see Fig.4). CycB:Cdk1, but not CycA.Cdk1, is regulated by inhibitory Cdk1-phosphorylation controlled by Wee1 and Cdc25. Both of the Tyr-modifying enzymes are regulated by CycB:Cdk1 itself as well as by CycA:Cdk1 and PP2A:B55. The activity of PP2A:B55 is controlled by the Greatwall-ENSA pathway which is activated by cyclin:Cdk1 dependent phosphorylation of Greatwall-kinase. The early mitotic (prophase) substrates could be phosphorylated by both cyclin:Cdk1 complexes and dephosphorylated by an unknown phosphatase. In contrast, the intermediate (prometaphase) and the late mitotic substrates are both dephosphorylated by PP2A:B55. While the intermediate substrate are phosphorylated by both cyclin:Cdk1 complexes, the phosphorylation of late substrates requires CycB:Cdk1 activity. We used nonlinear ordinary differential equations to describe the rate of change for protein concentrations in the network.

All the reactions are described by law of mass action kinetics except for the early mitotic substrates (pSearly) which is approximated by the Hill-equation. The rate constants are abbreviated by ‘k’ with subscripts referring to the type of reaction (a-activation, i-inhibition, p-phosphorylation and dp-dephosphorylation) and the substrate of the reaction. The numerical values of kinetic parameters are provided in the XPPAut ‘ode’ code that allows the users to reproduce Figure S14.

\# XPPAut code for simulation mitotic entry on Figure 4F and S14

CycA’ = kscyca - kdcyca*CycA
CycBT’ = kscycb*(1-CycBT)*CycBT
Wee1’ = (kawee1’ + kawee1*PP2AB55)^N*(1 - Wee1) - \ (kiwee1’*CycA + kiwee1*Cdk1)^N*Wee1
Cdc25P’ = (kaCdc25’*CycA + kaCdc25*Cdk1)^N*(1 - Cdc25P) - \ (kiCdc25’ + kiCdc25*PP2AB55)^N*Cdc25P
Cdk1’ = (kacdk1’ + (kacdk1 “-kacdk1 ‘)*Cdc25P)*(CycBT - Cdk1) - \ (kicdk1’ + (kicdk1 “-kicdk1 ‘)*Wee1)*Cdk1
Gwlp’ = (kaGwl’*CycA + kaGwl*Cdk1)*(1 - Gwlp) - kiGwl*PP2AB55*Gwlp
pENSAt’ = kpEnsa*Gwlp*(ENSAtot - pENSAt) - kdpEnsa*(B55tot - PP2AB55)
PP2AB55’ = (kdiss + kdpEnsa)*(B55tot - PP2AB55) - kass*PP2AB55*(pENSAt - (B55tot - PP2AB55))
pSearly’ = (kpearly’*CycA + kpearly*Cdk1)^M*(1 - pSearly) - kdpearly^M*pSearly
pSinter’ = (kpinter’*CycA + kpinter*Cdk1)*(1 - pSinter) - kdpinter*PP2AB55*pSinter
pSlate’ = (kplate’*CycA + kplate*Cdk1)*(1 - pSlate) - kdplate*PP2AB55*pSlate init CycA=0, CycBT=0.01, Wee1=1, Cdc25P=0, Cdk1=0, Gwlp=0, pENSAt=0,

PP2AB55=0.25, pSearly=0, pSinter=0, pSlate=0

\# Values of kinetic parameters
\# for Cyclin synthesis & degradation
p kscyca=0.02, kdcyca=0.02, kscycb=0.1
\# for Wee1 & Cdc25 activation &inactivation
p kawee1’=0.2, kawee1=8, kiwee1’=1, kiwee1=2, N=2
p kaCdc25’=1, kaCdc25=2, kiCdc25’=0.2, kiCdc25=8,
\# for Cdk1 activation & inactivation
p kacdk1’=0.01, kacdk1”=1, kicdk1’=0.01, kicdk1”=1
\# for Greatwall, ENSA & PP2A:B55
p kaGwl’=0.15, kaGwl=0.5, kiGwl=10
p ENSAtot=1, kpEnsa=6, kdpEnsa=3
p B55tot=0.25, kass=3600, kdiss=0.4
\# for phosphorylation & dephosphorylation of mitotic substrates
p kpearly’=1, kpearly=7, kdpearly=0.05, M=5
p kpinter’=0.1, kpinter=10, kdpinter=5
p kplate’=0, kplate=1, kdplate=100 @ xp=time, yp=CycA,xlo=0,xhi=100,ylo=0,yhi=1, total=100, meth=stiff @ nplot=8, yp=CycA, yp2=CycBT, yp3=Cdk1, yp4=pENSAt, yp5=PP2AB55, yp6=pSearly, yp7=pSinter, yp8=pSlate done

#### Statistical Analysis

All experiments included at least three independent biological repeats. Sample size per repeat varied between experiments and are indicated in the Figure Legends. Sample size was based on standard practise in cell biological assays and not specifically pre-estimated. P-values were calculated using an independent two sample t-test. Levels of significance are indicated by stars (* p<0.05, ** p<0.01, ***p<0.001). For all experiments, samples were not randomized and the investigators were not blinded to the group allocation during experiments and outcome assessment. No exclusion criteria were used and all collected data were used for statistical analysis.

## References

1. Dephoure, N. et al. A quantitative atlas of mitotic phosphorylation. Proc. Natl. Acad. Sci. USA 105, 10762–10767 (2008).

2. Daub, H. et al. Kinase-selective enrichment enables quantitative phosphoproteomics of the kinome across the cell cycle. Mol. Cell 31, 438–448 (2008).

3. Hégarat, N., Rata, S. & Hochegger, H. Bistability of mitotic entry and exit switches during open mitosis in mammalian cells. Bioessays 38, 627–643 (2016).

4. Fung, T. K. & Poon, R. Y. C. A roller coaster ride with the mitotic cyclins. Semin. Cell Dev. Biol. 16, 335–342 (2005).

5. Hochegger, H., Takeda, S. & Hunt, T. Cyclin-dependent kinases and cell-cycle transitions: does one fit all? Nat Rev Mol Cell Biol 9, 910–916 (2008).

6. Kalaszczynska, I. et al. Cyclin A Is Redundant in Fibroblasts but Essential in Hematopoieticand Embryonic Stem Cells. Cell 138, 352–365 (2009).

7. Fung, T. K., Ma, H. T. & Poon, R. Y. C. Specialized roles of the two mitotic cyclins in somatic cells: cyclin A as an activator of M phase-promoting factor. Mol. Biol. Cell 18, 1861–1873 (2007).

8. Gong, D. et al. Cyclin A2 regulates nuclear-envelope breakdown and the nuclear accumulation of cyclin B1. Curr. Biol. 17, 85–91 (2007).

9. Gong, D. & Ferrell, J. E. The roles of cyclin A2, B1, and B2 in early and late mitotic events. Mol. Biol. Cell 21, 3149–3161 (2010).

10. Deibler, R. W. & Kirschner, M. W. Quantitative Reconstitution of Mitotic CDK1 Activation in Somatic Cell Extracts. Mol. Cell 37, 753–767 (2010).

11. Gheghiani, L., Loew, D., Lombard, B., Mansfeld, J. & Gavet, O. PLK1 Activation in Late G2 Sets Up Commitment to Mitosis. CellReports 19, 2060–2073 (2017).

12. Vigneron, S. et al. Cyclin A-cdk1-Dependent Phosphorylation of Bora Is the Triggering Factor Promoting Mitotic Entry. Developmental Cell 45, 637–650.e7 (2018).

13. Burkard, M. E. et al. Chemical genetics reveals the requirement for Polo-like kinase 1 activity in positioning RhoA and triggering cytokinesis in human cells. Proc Natl Acad Sci USA 104, 4383–4388 (2007).

14. Katsuno, Y. et al. Cyclin A-Cdk1 regulates the origin firing program in mammalian cells. Proc. Natl. Acad. Sci. USA 106, 3184–3189 (2009).

15. Brandeis, M. et al. Cyclin B2-null mice develop normally and are fertile whereas cyclin B1-null mice die in utero. Proc Natl Acad Sci USA 95, 4344–4349 (1998).

16. Strauss, B. et al. Cyclin B1 is essential for mitosis in mouse embryos, and its nuclear export sets the time for mitosis. J. Cell Biol. 217, 179–193 (2018).

17. Chen, Q., Zhang, X., Jiang, Q., Clarke, P. R. & Zhang, C. Cyclin B1 is localized to unattached kinetochores and contributes to efficient microtubule attachment and proper chromo-some alignment during mitosis. Cell Res 18, 268–280 (2008).

18. Castilho, P. V., Williams, B. C., Mochida, S., Zhao, Y. & Goldberg, M. L. The M phase kinase Greatwall (Gwl) promotes inactivation of PP2A/B55delta, a phosphatase directed against CDK phosphosites. Mol. Biol. Cell 20, 4777–4789 (2009).

19. Mochida, S., Maslen, S. L., Skehel, M. & Hunt, T. Greatwall phosphorylates an inhibitor of protein phosphatase 2A that is essential for mitosis. Science 330, 1670–1673 (2010).

20. Gharbi-Ayachi, A. et al. The substrate of Greatwall kinase, Arpp19, controls mitosis by inhibiting protein phosphatase 2A. Science 330, 1673–1677 (2010).

21. Burgess, A. et al. Loss of human Greatwall results in G2 arrest and multiple mitotic defects due to deregulation of the cyclin B-Cdc2/PP2A balance. Proc. Natl. Acad. Sci. USA 107, 12564–12569 (2010).

22. Alvarez-Fernández, M. et al. Greatwall is essential to prevent mitotic collapse after nuclear envelope breakdown in mammals. Proc. Natl. Acad. Sci. USA 110, 17374–17379 (2013).

23. Cundell, M. J. et al. A PP2A-B55 recognition signal controls substrate dephosphorylation kinetics during mitotic exit. J. Cell Biol. 214, jcb.201606033–554 (2016).

24. Natsume, T., Kiyomitsu, T., Saga, Y. & Kanemaki, M. T. Rapid Protein Depletion in Human Cells by Auxin-Inducible Degron Tagging with Short Homology Donors. CellReports 15, 210–218 (2016).

25. Chung, H. K. et al. Tunable and reversible drug control of protein production via a self-excising degron. Nature Chemical Biology 11, 713–720 (2015).

26. Lemmens, B. et al. DNA Replication Determines Timing of Mitosis by Restricting CDK1 and PLK1 Activation. Mol. Cell 71, 117–128.e3 (2018).

27. Zerjatke, T. et al. Quantitative Cell Cycle Analysis Based on an Endogenous All-in-One Reporter for Cell Tracking and Classification. Cell Reports 19, 1953–1966 (2017).

28. Sackton, K. L. et al. Synergistic blockade of mitotic exit by two chemical inhibitors of the APC/C. Nature 514, 646–649 (2014).

29. Cuylen, S. et al. Ki-67 acts as a biological surfactant to disperse mitotic chromosomes. Nature 535, 308–312 (2016).

30. Blethrow, J. D., Glavy, J. S., Morgan, D. O. & Shokat, K. M. Covalent capture of kinase-specific phosphopeptides reveals Cdk1-cyclin B substrates. Proc. Natl. Acad. Sci. USA 105, 1442–1447 (2008).

31. Tsukahara, T., Tanno, Y. & Watanabe, Y. Phosphorylation of the CPC by Cdk1 promotes chromosome bi-orientation. Nature 467, 719–723 (2010).

32. Goto, H. et al. Complex formation of Plk1 and INCENP required for metaphase–anaphase transition. Nature Publishing Group 8, 180–187 (2005).

33. Cundell, M. J. et al. The BEG (PP2A-B55/ENSA/Greatwall) pathway ensures cytokinesis follows chromosome separation. Mol. Cell 52, 393–405 (2013).

34. Ly, T. et al. Proteomic analysis of cell cycle progression in asynchronous cultures, including mitotic subphases, using PRIMMUS. eLife 6, 843 (2017).

35. Bouchoux, C. & Uhlmann, F. A Quantitative Model for Ordered Cdk Substrate Dephosphorylation during Mitotic Exit. Cell 147, 803–814 (2011).

36. Rata, S. et al. Two Interlinked Bistable Switches Govern Mitotic Control in Mammalian Cells. Curr. Biol. 0, (2018).

## References

1. F. A. Ran et al., Genome engineering using the CRISPR-Cas9 system. Nat. Protoc. 8, 2281–2308 (2013).

2. T. Natsume, T. Kiyomitsu, Y. Saga, M. T. Kanemaki, Rapid Protein Depletion in Human Cells by Auxin-Inducible Degron Tagging with Short Homology Donors. CellReports. 15, 210–218 (2016).

3. T. Zerjatke et al., Quantitative Cell Cycle Analysis Based on an Endogenous All-in-One Reporter for Cell Tracking and Classification. Cell Reports. 19, 1953–1966 (2017).

4. K. J. Livak, T. D. Schmittgen, Analysis of relative gene expression data using real-time quantitative PCR and the 2(-Delta Delta C(T)) Method. Methods. 25, 402–408 (2001).

5. S.-I. Hiraga et al., Human RIF1 and protein phosphatase 1 stimulate DNA replication origin licensing but suppress origin activation. EMBO Rep. 18, 403–419 (2017).

6. J. Cox, M. Mann, MaxQuant enables high peptide identification rates, individualized p.p.b.-range mass accuracies and proteome-wide protein quantification. Nat Biotechnol. 26, 1367–1372 (2008).

7. J. Cox et al., Andromeda: a peptide search engine integrated into the MaxQuant environment. J Proteome Res. 10, 1794–1805 (2011).

8. S. Rata et al., Two Interlinked Bistable Switches Govern Mitotic Control in Mammalian Cells. Curr. Biol. 0 (2018), doi:10.1016/j.cub.2018.09.059.

